# Bio-macromolecular assembly machine for human limb lengthening

**DOI:** 10.1101/2024.08.15.608028

**Authors:** Chang Xie, Wenyue Li, Xudong Yao, Boxuan Wu, Jinghua Fang, Renwei Mao, Yiyang Yan, Hongxu Meng, Yan Wu, Xianzhu Zhang, Wangping Duan, Xuesong Dai, Xiaozhao Wang, Hongwei Ouyang

**Author notes:** Corresponding Author: Hongwei Ouyang, Ph.D. Department of Sports Medicine of the Second Affiliated Hospital, and Liangzhu Laboratory, Zhejiang University School of Medicine, Hangzhou, Zhejiang, China; Dr. Li Dak Sum & Yip Yio Chin Center for Stem Cells and Regenerative Medicine, Zhejiang University School of Medicine, Hangzhou, China. Corresponding Author: Xiaozhao Wang, Ph. D. Liangzhu Laboratory, Zhejiang University School of Medicine, Hangzhou, Zhejiang, China; Dr. Li Dak Sum & Yip Yio Chin Center for Stem Cells and Regenerative Medicine, Zhejiang University School of Medicine, Hangzhou, China. The authors contribute equally to this work (Chang Xie, Wenyue Li).

## Abstract

Growth plate (GP) is the critical cartilaginous structure for longitudinal bone growth. Herein, employing high-resolution analytical techniques, we explore the intricate mechanisms that govern the polarized mineralization patterns within GP. The GP-epiphysis interface displays a sharp transition in tissue modulus, acting as a “protective shell” for the underlying GP, whereas the GP-metaphysis interface exhibits a gradual modulus increase, enabling efficient load redistribution to metaphysis. The unique mechanical environments at these two interfaces contribute to polarized CaP mineralization patterns, which are regulated by a complex protein-based molecular machinery. The mineralization inhibitors SPP1 and AHSG enriched at the GP-epiphysis interface could act as a line of defence against mineralization. When these two proteins coexisted with the mineralization-promoting ENPP1 and ALPL at the GP-metaphysis interface, a sequential event of precise nucleation and programmable assembly of CaP minerals occur, forming “mineralization waves” to guide bone elongation. By replicating such specific macromolecular environment at GP-metaphysis interface, a hypertable amorphous calcium phosphate (ACP) phase is well-retained *in vitro*, demonstrating the possibility for precise and gentle control of ACP-hydroxyapatite (HAp) transformation under physiological conditions. Our study defines a novel concept of “mineralization waves” that govern the velocity and amplitude of GP-guided mineralization process.

## Introduction

Organisms possess the ability to regulate the mineral assemblies for maintaining various physiological functions(*1*, *2*). Calcium phosphate (CaP), as the primary component of hard tissues like tooth enamel and bone in vertebrates, has been the subject of extensive research due to its controllable assembly(*3*, *4*). Current studies have focused on controlling CaP assembly through ions(*5*, *6*), peptides(*7*, *8*), proteins(*9*), DNA(*10*) and RNA(*11*). However, the complex phase transition processes involved in non-classical nucleation render CaP assembly uncontrollable. Despite the significant progress made in understanding the mineralization of hard tissues, the material science of soft tissue mineralization and ossification, particularly in growth plate (GP) responsible for long bone growth remains unexplored. GPs are the cartilaginous structures located at the ends of long bones and critical regions responsible for longitudinal bone growth and development in humans. The developmental GP-bone interfaces serve as an ideal template for studying human mineral growth because of their unique polarized mineralization pattern. Following the formation of the secondary ossification center, GPs are situated between epiphysis and metaphysis(*12*) (**Figure S1**). Long bone growth occurs through the GP mineralizing into the metaphysis via endochondral ossification(*13*, *14*). In contrast to the actively mineralized GP-metaphysis interface, the GP cartilage does not undergo continuous mineralizing into bone tissue at the GP-epiphysis interface(*15*). This phenomenon maintains the polarized nature of bone elongation, progressing from the GP towards the metaphysis. Previous studies have extensively revealed the unique contribution of disparate cell types within the GP in guiding long-term bone elongation(*16*, *17*). However, the polarized mineralization patterns of GPs have rarely been explored from a materials science perspective, especially in human samples.

GPs are located in a unique hard-soft-hard mechanical environment between the epiphysis and metaphysis. Under physiological conditions, bone tissue exhibits remarkable adaptation to the mechanical environment, dynamically responding to biomechanical stress and generating mineralized structures and compositions optimized for the mechanical response(*18*). Understanding the interplay of localized mechanical responses as well as the corresponding microstructural and compositional transitions of the two key cartilage-bone interfaces can help explain the polarized biomineralization pattern in GPs. Moreover, the process of biological mineralization is a finely regulated process orchestrated by cells, involving the transient stabilization of amorphous minerals controlled by biomacromolecules(*2*), nucleation(*19*), crystal growth and three-dimensional assembly within the extracellular matrix (ECM)(*20*). Mineralization inhibitors at the GP-epiphysis interface effectively deter the precipitation of abundant calcium and phosphate ions within the extracellular matrix(*9*, *21*). While at the GP-metaphysis region, the inhibitory effect of mineralization is neutralized by biomineralization promoters like enzymes(*22*, *23*), allowing calcium and phosphate ions to form mineral precursors for further crystalization(*4*, *24*, *25*). These processes are believed to be controlled by a macromolecular machine comprising inhibitors and promotors that regulate mineralization dynamics at the GP interfaces. A comprehensive understanding of the underlying biomechanical, biochemical, structural and biomolecular mechanisms involved in the polarized mineralization patterns at the GP-bone interfaces is of great significance in fields such as controlled mineral assembly, bone growth and development studies, and bone tissue engineering.

Herein, we collected human GP samples (phalange GPs: 0-5 years old; tibia GPs: 6-14 years old) (**Table S1**) to reveal the polarized bone lengthening process, including mechanical microenvironment, microstructural and compositional transformation, as well as nano-scale crystal assembly of the key transitional GP-epiphysis and GP-metaphysis regions by multiple high-resolution imaging technologies (**Figure S2**). Our results confirmed the existence of a mineral precursor phase which shared high similarities with amorphous calcium phosphate (ACP) during GP biomineralization. Proteomics revealed a series of proteins that maintain the mineralization inhibition region at the GP-epiphysis interface, stabilize the ACP-like precursor, and regulate the hierarchical mineralization process from GP to metaphysis by acting as a biomolecular machine. Based on these findings, we successfully stabilized ACP for at least 35 days at 37 °C and manipulated the ACP to hydroxyapatite (HAp) transformation *in vitro*, proposing a new mechanism termed “mineralization waves” to describe the dynamic process of mineral assembly and transformation observed in GP biomineralization.

## Results

### Mechanical characteristics of the GP-epiphysis/metaphysis interface

The *in-situ* biomechanical performance of tibia GP tissue, including epiphysis and metaphysis, under physiological loading was evaluated at high resolutions (5.64 μm per voxel) by synchrotron X-ray microscope (XRM) in combination with mechanical loading (**Figure 1A**). The tested sample underwent heterogenous deformation after compressive loading (**Figure S3A**). We reconstructed the 3D structure before and after loading by Micro-computed tomography (micro-CT) and conducted Digital Volume Correlation (DVC) to quantify local displacement and strain(*26*) (**Figure S3B**). The strain in the mineralized region spanning from the epiphysis to the metaphysis were revealed, and the GP strain could be examined via two interfaces of the GP, while internal deformation of the GP was indiscernible because of deficient voxel differentiation. The GP-metaphysis interface showed more significant load-induced displacements compared to the GP-epiphysis interface upon loading (**Figure 1B, Movie S1-3**). A sharp increase in local strain was detected from the GP to epiphysis, whereas the transition from the GP to the metaphysis exhibited a more gradual rise in strain level (**Figure 1C**), indicating that the former interface undergoes a sudden change in mechanical properties while the latter exhibits a gradient transition.

**Figure 1:**
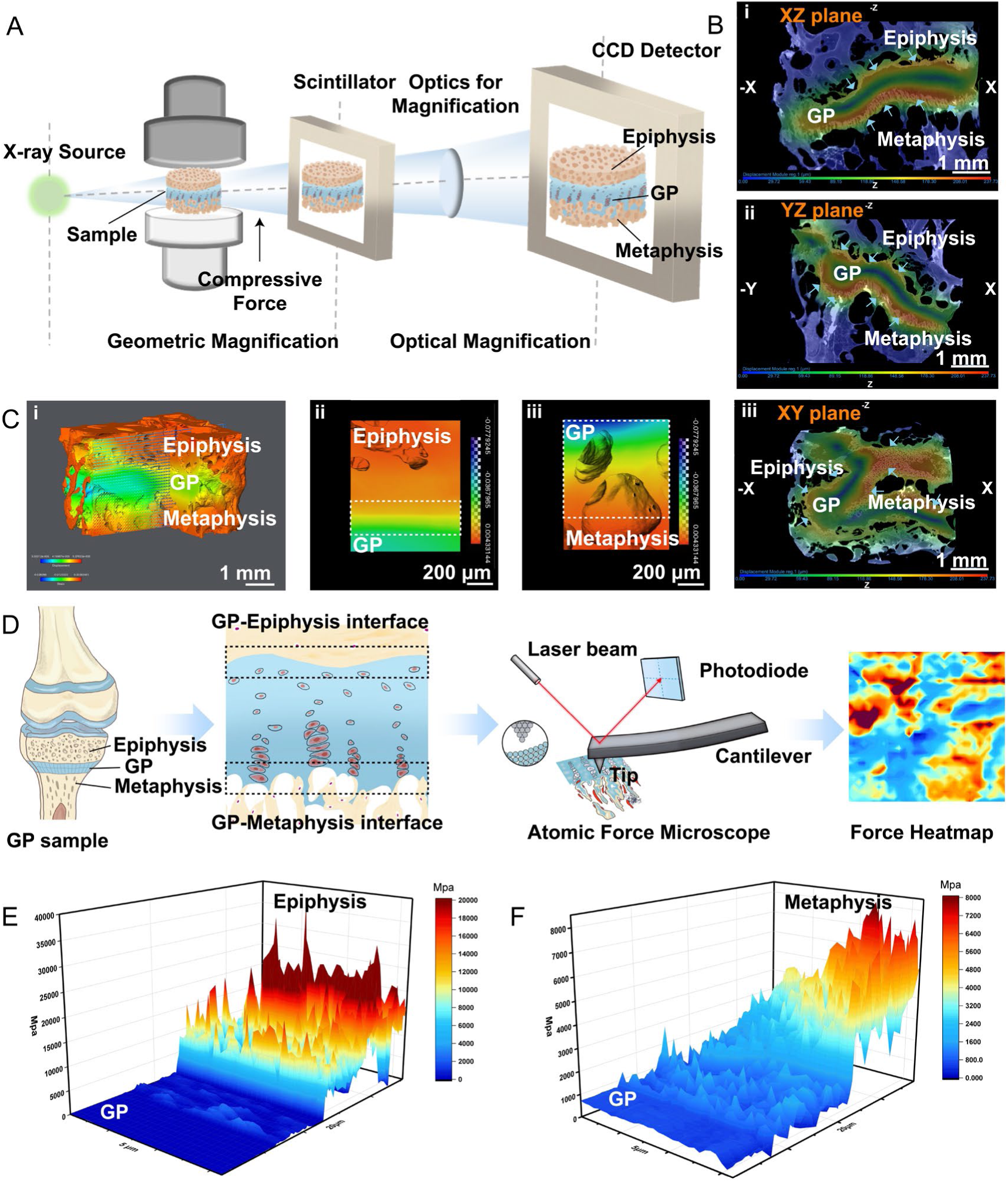
The mechanical response and properties of the growth plate. A) Schematic illustration of the in situ micro-CT imaging of the whole GP sample via synchrotron XRM; B) The displacement magnitude images via DVC processing of GP tissue under compression in the (i) XZ, (ii) YZ and (iii) XY plane, The arrows point to the interfaces with significant displacement changes, scale bar = 1mm; C) Views of (i) 3D strain distribution of GP tissue when compression is applied, the arrows represent the displacement direction, scale bar = 1mm, and expanded views of the (ii) GP-epiphysis interface, scale bar = 200 μm, and (iii) GP-metaphysis interface, scale bar = 200μm, processed by DVC; D) Schematic of AFM analysis of the GP-Epiphysis and GP-metaphysis interface; E-F) The 3D modulus distribution map of all selected areas from GP to epiphysis/metaphysis.

Next, we evaluated the modulus transition from the GP to the epiphysis/metaphysis using atomic force microscopy (AFM) (**Figure 1D**). The tissue modulus increased sharply from 130.70 ± 36.56 MPa in the resting zone of the GP to 11.92 ± 6.60 GPa in the epiphysis (**Figure 1E, Figure S4A-B**). In contrast, the GP-metaphysis interface demonstrated a relatively progressive increment from 416.20 ± 107.18 MPa in the hypertrophic zone of the GP to 3.22 ± 1.53 GPa in the metaphysis (**Figure 1F, Figure S4C-D**). The distinct modulus transition patterns were consistent with the differential mechanical responses at the GP-epiphysis and GP-metaphysis interface. The epiphysis, with its remarkable stiffness, likely acts as a “protective shell” and prevents catastrophic damage to the resting zone in GPs upon impact, thus ensuring the survival of the stem cell reservoir and facilitating bone development(*27*). The stiffness gradient from the GP to the metaphysis facilitates load redistribution, which has been reported to be beneficial for biomineralization during bone lengthening(*28*, *29*).

### Distinct CaP mineralization patterns at two interfaces

During long bone lengthening, bone tissue dynamically responds to mechanical stress and optimizes its structures and compositions(*18*, *30*). We observed the structural and compositional transition at the soft-hard interfaces, including the GP-epiphysis and GP-metaphysis interface (**Figure 2A-G; Figure S5A**), using scanning electron microscopy (SEM), density dependent color-SEM (DDC-SEM), focused ion beam-SEM (FIB-SEM) and energy-dispersive X-ray spectroscopy (EDX). At the GP-epiphysis interface, the porous and loose GP cartilage in the resting zone underwent a sharp transformation into dense and mature epiphysis tissue (**Figure 2H; Figure S5B**). Meanwhile, at the GP-metaphysis interface, densely packed collagen fibers in calcified zones gradually mineralized into bone tissue, with minerals transitioning from immature spherical minerals to crystal platelets (**Figure 2I; Figure S5C**). FIB-SEM also showed extrafibrillar spherical minerals at the mineralization front, which fused across the interfibrillar space and permeated the collagen fibrils to transform into a fully mineralized metaphysis region (**Figure S6**). EDX line scans and mapping of calcium (Ca) and phosphorus (P) distribution further highlighted the differences between the sharp elemental transformation across the GP-epiphysis interface (**Figure 2J, Figure S7A**) and the gradual transition across the GP-metaphysis interface occurring over a longer distance (**Figure 2K, Figure S7B**). Moreover, the minerals in the epiphysis region exhibited Ca/P ratio analogous to mature HAp crystals, and this value remained consistent throughout the epiphysis (zone 1, 1.763±0.067; zone 2, 1.693±0.050; zone 3, 1.660±0.043) (**Figure S5D, S5F**). In contrast, the Ca/P ratios of the metaphysis tissue near the mineralization front (zone 1, 2.205±0.190; zone 2, 1.930±0.072) were significantly higher than those far from the front (zone 3, 1.612±0.085) (**Figure S5E, S5G**), indicating the possible existence of an amorphous phase and the gradual maturation of mineral deposits from amorphous to crystalline minerals across this interface(*31*).

**Figure 2:**
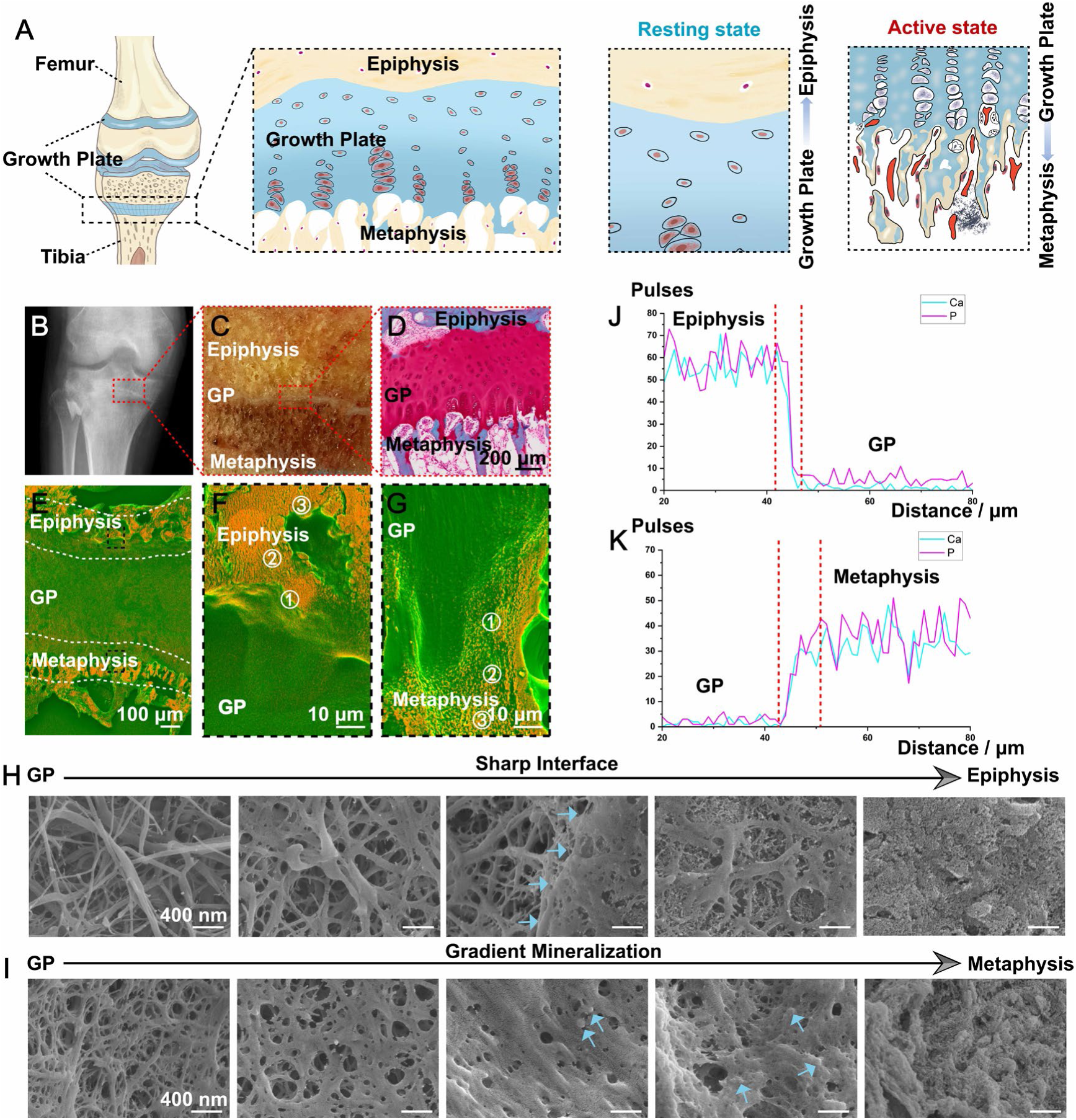
Microstructural and compositional transformation of human GP interfaces. A) The schematic illustration of human tibia growth plate, with resting GP-epiphysis interface and active GP-metaphysis interface; B) X-ray image of human knee joint with GP; C) Representative photograph of GP, epiphysis and metaphysis in tibia; D) SO staining of GP, scale bar = 200μm; E) The DDC-SEM micrograph of GP with epiphysis and metaphysis tissue, scale bar = 100μm; F-G) Enlarged DDC-SEM micrographs of the GP-epiphysis interface and GP-metaphysis interface, scale bar = 10μm; H, I) Representative enlarged SEM images from the GP to epiphysis and the GP to metaphysis, blue arrows pointing to the sharp transformation from the GP to epiphysis in H) and mineral particles in I), scale bar = 400nm; J, K) The EDX line scanning from J) the epiphysis to GP and K) the GP to metaphysis, showing Ca and P distribution across the interfaces, the red dotted line marked the interface region.

We performed stimulated Raman scattering (SRS) imaging to further evaluate the mineral phase and the spatial component distribution such as glycosaminoglycans (GAGs), protein, lipid (**Figure S8**) and especially minerals across the two interfaces. The abrupt increase of PO_4_^3-^*v*_1_ symmetric stretching peak at 960 cm^-1^ indicated that the GP-epiphysis interface underwent a sharp transition from non-mineralized GP cartilage to fully mineralized epiphysis occupied by carbonated crystalline HAp (**Figure 3A-3B, Figure S9A**). In contrast, the GP-metaphysis interface displayed a distinct pattern: a noticeable shift in the broad peak of the PO_4_^3-^*v*_1_ band occurred at 950 cm^−1^ and 955 cm^−1^, which finally transformed into the narrow peak at 960 cm^-1^ (**Figure 3C, Figure S9B**). The appearance of the peak at 955 cm^−1^ may be caused by overlapping peaks of ACP at 950 cm^−1^ and HAp at 960 cm^−1^(*32*). Additionally, SRS mapping validated the presence of a region enriched with ACP at the frontier of the GP-metaphysis interface, which subsequently mineralized into HAP (**Figure 3D**). These results align with the observed spherical structures in SEM and high Ca/P ratios at the GP-metaphysis interface, suggesting an ACP-HAp transformation during mineralization from the GP into the metaphysis(*32–34*). Closer inspection of the mineral spatial distribution showed an increase in HAp crystallinity, substituted carbonate content and mineral-to-collagen ratio, as well as a decrease CO_3_^2–^/PO_4_ ^3–^ ratio across the GP-epiphysis interface (**Figure 3E-3G**). In contrast, a more gradual trend was observed from the GP to the metaphysis (**Figure 3H-3J**). Collectively, these results revealed two different mineralization distribution patterns at the GP-epiphysis interfaces (sharp HAP distribution) and the GP-metaphysis interface (gradual ACP-HAp transformation).

**Figure 3:**
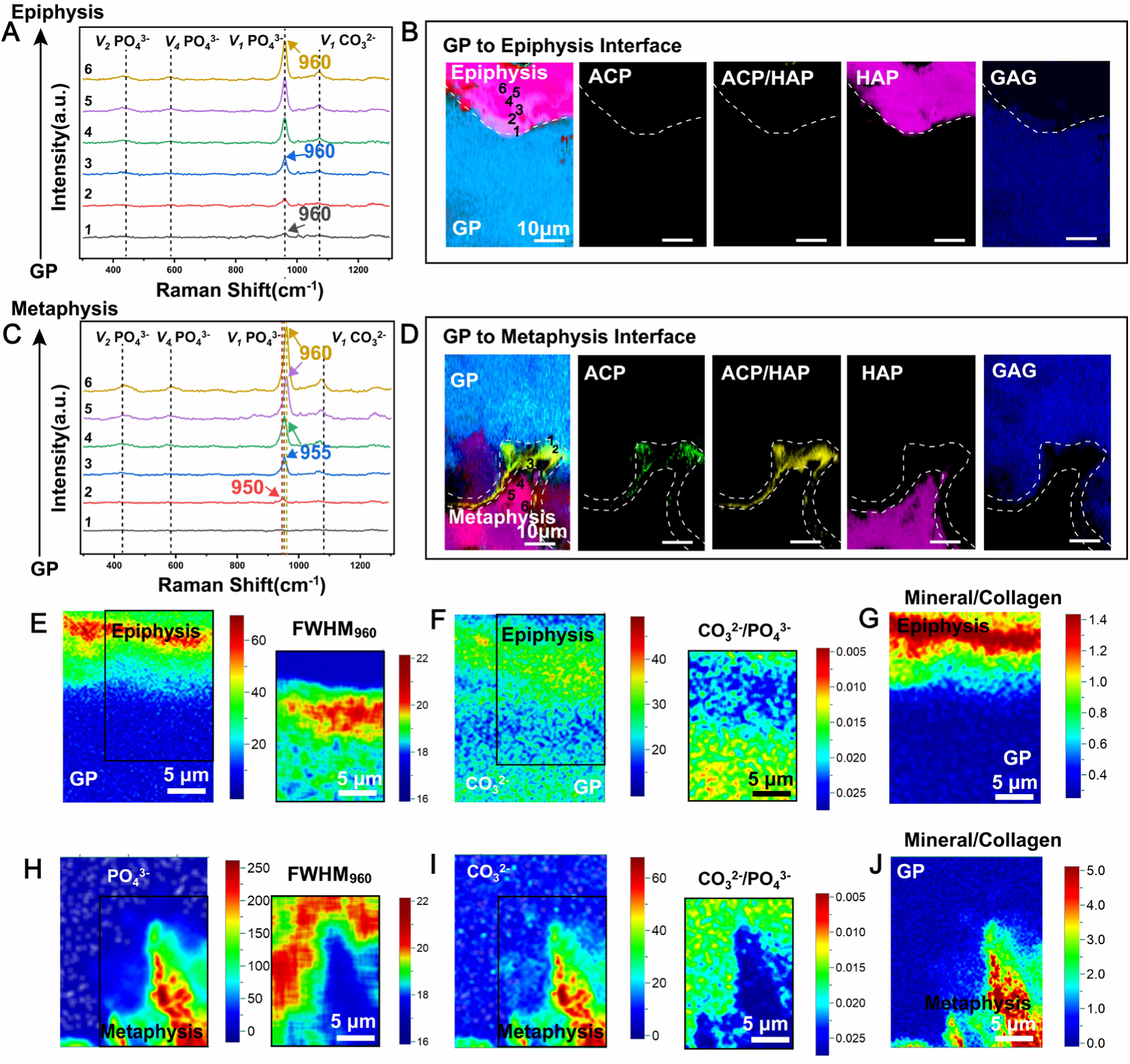
The chemical distribution maps of the GP-epiphysis interface and GP-metaphysis interface revealed by Raman microscopic detection technology. A, C) Raman spectra collected in the 300∼1300 cm^-1^ from GP to A) epiphysis and C) metaphysis at different area (zone 1-6) in B) and D), The information of marked Raman shifts: PO_4_^3-^*v*_1_ band at 950∼960 cm^-1^, PO_4_^3-^*v*_2_ band at 430∼433 cm^-1^ and PO_4_^3-^*v*_4_ band at 573∼590 cm^-1^ and CO_3_^2–^*v*_1_ band at 1071 cm^−1^; B, D) The maps of ACP (950 cm^-1^), ACP/HAp interfering peaks (955 cm^-1^), HAp (960 cm^-1^) and GAG (1410 cm^-1^) in B) GP-epiphysis interface and D) GP-metaphysis interface by SRS imaging, scale bar = 10μm; E, H) Raman peak intensity maps of PO ^3-^*v* band at 960 cm^-1^ and corresponding maps of full-widths at half-maximum (FWHM) of the peak at 960 cm^-1^ at E) GP-epiphysis interface and H) GP-metaphysis interface, scale bar = 5μm; F, I) The maps of CO_3_^2-^v_1_ band at 1071cm^-1^ and CO_3_ ^2-^/ PO_4_ ^3-^ratio at F) GP-epiphysis interface and I) GP-metaphysis interface, revealing the ionic substitution of minerals in both interfaces, scale bar =5μm; G, J) The mineral to collagen maps (960 cm^-1^ to 1595∼1700 cm^-1^) of G) GP-epiphysis interface and J) GP-metaphysis interface, revealing relative content and distribution of HAp, scale bar = 5μm.

### Disparate chemical environments contribute to distinct CaP transformation patterns at two interfaces

To uncover the nature of transitioning minerals at the two interfaces, we investigated nano-scale CaP mineral assembly by cryo-transmission electron microscopy (cryo-TEM) and selected area electron diffraction (SAED). The nanocrystals with diameters ranging from 5-10 nm appeared at the frontier of the GP-epiphysis interface (**Figure 4Ai, 4Bi**), which gradually transformed into bulk platelets (**Figure 4A-4C, Figure S10-S11, S12A, S13**). Differently, at the mineralization front of the GP-metaphysis interface, amorphous clusters with total dimensions of 150-200 nm, consisting of nanometer-sized building blocks, strands of spherical units, and nodules, were observed (**Figure 4Di, 4Ei, Figure S14A, Figure S15A, Figure S16A-B and Figure S17A**)(*5*, *19*). These structures corresponded to the spherical structures and amorphous phase shown in Figure 2 and Figure 3, respectively. As mineralization progressed, the ACP-like clusters mineralized into poorly crystalline nanoparticles with diameters of 5-10 nm, which continued to grow by oriented particle attachment, a process frequently found in the early stages of crystallization in the non-classical mineralization pathway(*19*, *20*, *35*)(**Figure 4Dii, 4Eii, Figure 4F, Figure S12B, Figure S14B-D, Figure S15B-C**). These nanoparticles eventually transformed into single bulk crystals. The crystal platelets then aggregated into mineral spherulites (**Figure 4Diii, Figure S14E, Figure S15D-E, Figure S16C-D**), which spread across the collagen fibrils and grew larger until complete mineralization of the fibrils (**Figure 4Div and 4Eiv, Figure S14F-G, Figure S15F, Figure S16E-G**). Similar mineral assembly processes at the GP-epiphysis and GP-metaphysis interface were also observed in GP samples of human phalange tissue (**Figure S13, S16-S17**).

**Figure 4:**
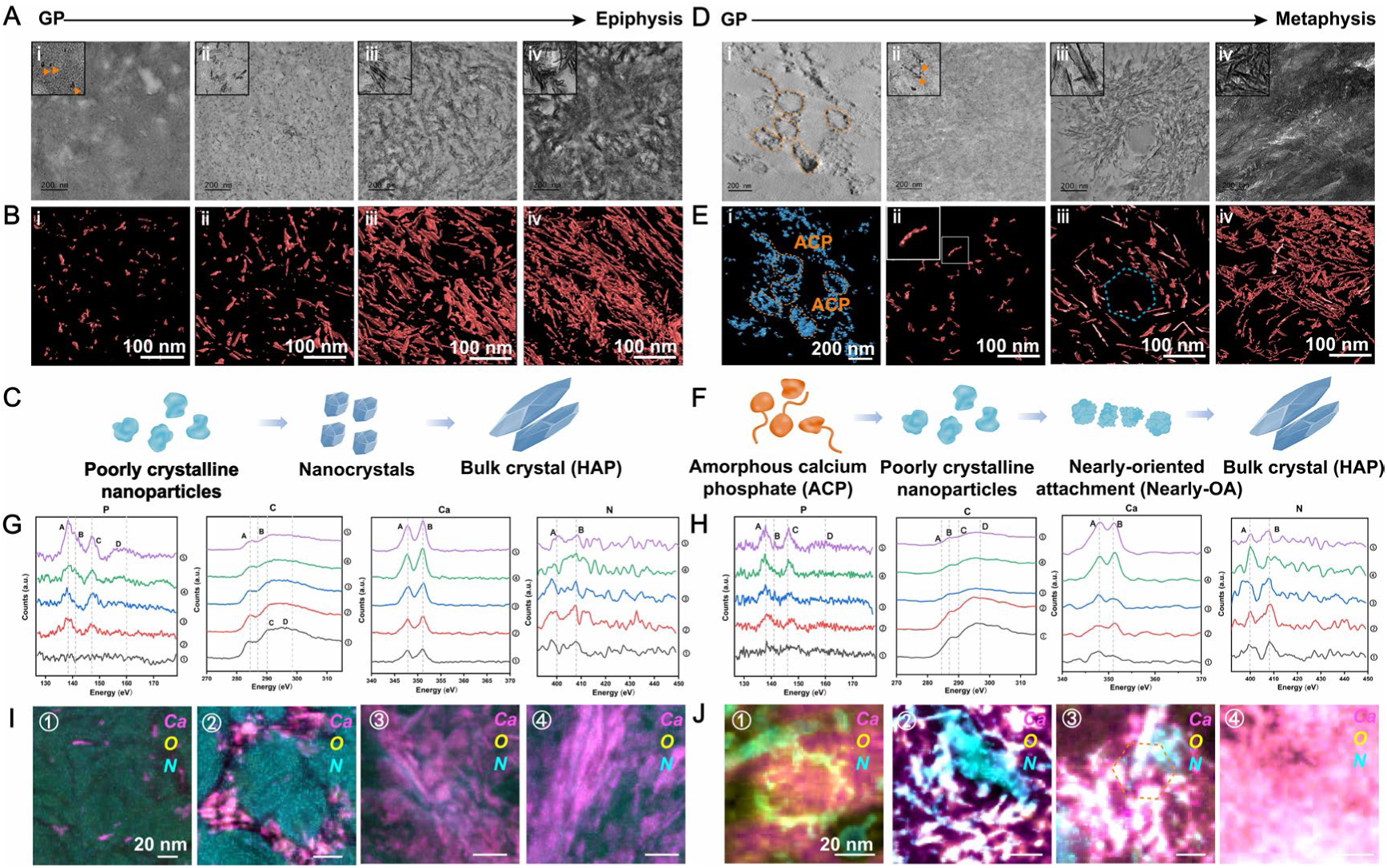
The CaP transformation pattern from the GP to epiphysis/metaphysis. A-B) Representative cryo-TEM images from GP to epiphysis and reconstructed tomographic pictures of cryo-TEM images, scale bar = 100 nm; C) Schematic illustration of crystal assembly process at the GP-epiphysis interface; D-E) Representative cryo-TEM images from GP to metaphysis and reconstructed tomographic pictures of cryo-TEM images, scale bar = 100 nm; F) Schematic illustration of the crystal assembly process at the GP-metaphysis interface; G-H) EELS spectra taken at P L_23_-edge, C K-edge, Ca L_23_-edge, N K-edge and O K-edge collected from different mineral particles in zone 1-4 in I) and J); I-J) EELS maps corresponding to the spatial distribution of Ca in magenta (collected at Ca L_23_-edge), O in yellow (collected at O K-edge) and N in cyan (collected at N K-edge) of selected HAADF-STEM images in Figure S18, from I) GP-epiphysis and J) metaphysis, scale bar = 20 nm.

To further elucidate the local chemical microenvironment of each mineral phase at two GP interfaces, an analysis using high-angle annular dark-field-scanning transmission electron microscopy (HAADF-STEM) equipped with electron energy loss spectroscopy (EELS) was performed **(Table S2)**. The signal intensity of calcium (Ca L_23_ edge) and phosphate (P L_23_ edge) increased across both interfaces, showing the increasing inorganic mineral content. Simultaneously, the carbonate and nitrogen signals were more intense at the front of both soft-hard interfaces and became weaker as the mineral matures (**Figure 4G-4J, Figure S18-S19**). Moreover, carbonyl groups (peak B) and nitrogen (peak B) signals were interspersed in ACP-like structures and poorly crystalline minerals at the GP-metaphysis interface (**Figure 4H, 4J**), indicating the presence of macromolecules that stabilize the amorphous phase and regulate the amorphous-to-crystalline transformation during GP mineralization(*31*).

### The biomolecule-based regulatory mechanism of GP-guided polarized long bone elongation

Next, we performed liquid chromatography-tandem mass spectrometry (LC-MS/MS) to reveal regulatory proteins at the interfaces **(Figure 5A, Figure S20)**. Among the 4,678 proteins detected, secreted phosphoprotein 1 (SPP1) and Alpha 2-HS glycoprotein (AHSG) were significantly enriched at the GP-epiphysis interface compared to the GP (**Figure 5B**), which was further confirmed by immunofluorescence staining (**Figure 5D, Figure S21A-B**). SPP1 and AHSG are able to chelate Ca ions to inhibit HAp nucleation and growth(*36*), leading to the suppression of mineralization at the GP-epiphysis interface. Enrichment of SPP1 and AHSG was also found at the GP-metaphysis interface compared to GP tissue (**Figure 5C, 5E and Figure S21C-D**), In this case, enrichment likely contributes to the transient stabilization of ACP and poorly crystalline minerals, preventing excessively rapidly HAp formation during GP mineralization. Besides, Ecto-nucleotide pyrophosphatase phosphodiesterase 1 (ENPP1) and alkaline phosphatase (ALPL) are highly expressed at the front of GP-metaphysis interface (**Figure 5C, 5E and Figure S21E-S21F**). ENPP1 can hydrolyze adenosine triphosphate (ATP), generating adenosine monophosphate (AMP) and pyrophosphate (PPi), which is then hydrolyzed by ALPL to inorganic phosphate (Pi), thus providing phosphate sources to facilitate the mineralization process at the GP-metaphysis interface(*37*). In coordination with calcium ions provided by SPP1 and AHSG, proteins expressed at the GP-metaphysis interface function as a biomolecular machine and provide a biomineralization-promoting condition for rapid bone lengthening.

**Figure 5:**
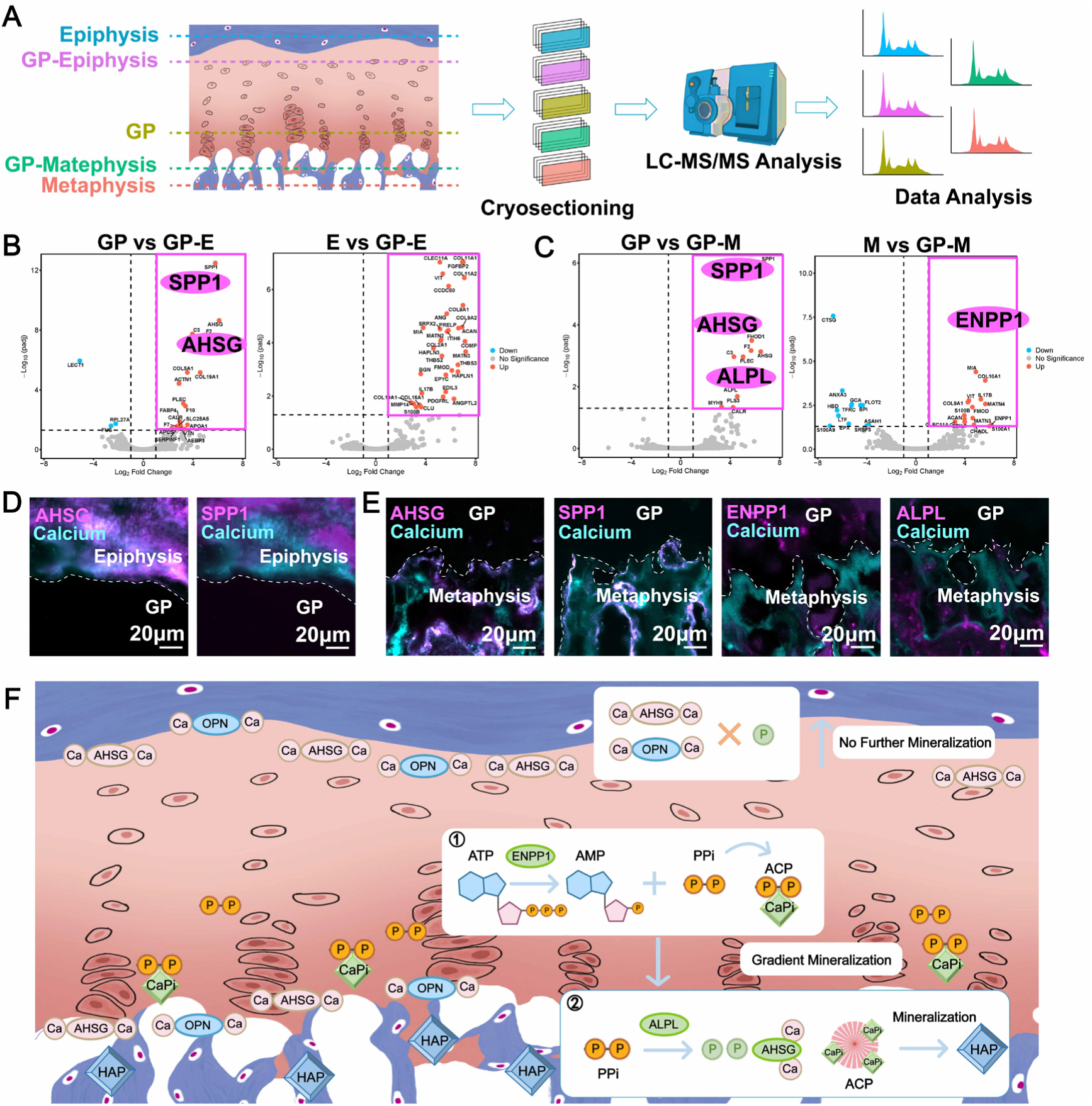
Regulatory mechanism of polarized GP mineralization revealed by proteomics. A) Experimental procedure for LC-MS/MS. The GP samples from 3 biological replicates were divided into five groups: epiphysis (E), GP-epiphysis (GP-E), GP, GP-metaphysis (GP-M) and metaphysis (M) tissue; B-C) Volcano plot showing proteins with differential expression between interfaces and GP tissue/bone tissue (epiphysis and metaphysis) (dotted line showing p-value < 0.05 and two-fold change cut-offs); SPP1, AHSG, ENPP1 and ALPL have been highlighted in the volcano plot; D) Representative images of GP samples immunostained for AHSG (magenta) /calcium (cyan) and SPP1(magenta) /calcium (cyan) at the GP-epiphysis interface; E) Representative images of GP samples immunostained for AHSG/calcium, SPP1/calcium, ENPP1 /calcium and ALPL/calcium at the GP-metaphysis interface, scale bar = 20 μm; F) Summary diagram of the macromolecule-based regulatory mechanism of GP-guided polarized long bone elongation.

In addition to the biomacromolecules, ions such as iron (Fe^2+^) and strontium (Sr^2+^), detected by FIB-SEM equipped with time-of-flight secondary ion mass spectrometry (TOF-SIMS), were only enriched in a 10 μm-wide region at the front of the GP-epiphysis interface compared to the GP-metaphysis interface (**Figure S22**). This ion distribution map, which colocalized with the stiff “protective shell” in Figure 1, may also play a role in inhibiting crystal growth(*5*, *38*) and maintaining the inhibition state at the GP-epiphysis interface.

Our findings outline the macromolecule-based regulatory mechanism that governs GP-guided polarized long bone elongation (**Figure 5F**). At the GP-epiphysis interface, the abundant calcium and phosphate ions in the ECM are prevented from precipitation by biomineralization inhibitors like SPP1 and AHSG, which form an additional line of defence against mineralization. While at the GP-metaphysis interface, the key region of GP mineralization, the inhibitory effect of SPP1 and AHSG towards calcium phosphate mineralization is abolished by enzymatic activity of the resident cells(*24*), thus facilitating the ECM mineralization. PPi produced by hydrolyzation of ATP by ENPP1 at the beginning of GP mineralization could maintain an extracellular bioreservoir of phosphate. As the mineralization progresses, PPi is actively enzymatically cleaved by ALPL into Pi. The enzymatic activity of ALPL on PPi not only abolishes the inhibitory effect of biomineralization but also generate phosphate ions to promote mineralization. PPi or Pi-packed granules chelate calcium ions and form the disordered calcium phosphate as mineral precursors and promote intrafibrillar mineralization of bone tissue.

We next attempted to manipulate mineralization process, including maintaining a stable ACP phase and controlling ACP-HAp transformation *in vitro,* by mimicking the macromolecular machine. ATP was added to provide phosphorous source in this system. After sufficient reaction of ATP and ENPP1, ALPL was added to the solution to produce Pi by hydrolyzation of PPi. ASHG was added to the CaCl_2_ solution to form calciprotein particles and regulate calcium-phosphate deposition. Subsequently, the two solutions were mixed to produce calcium phosphate precursors (**Figure 6A**). After 5min of reaction at 37 °C, clusters with obvious contrast were observed by cryo-TEM, which revealed spherical amorphous particles with a diameter of 50-150 nm after 30min and exhibited typical ACP morphologies(*39*) after 2h (**Figure 6B**). SAED and EDS mapping further confirmed the non-crystalline state of the prepared ACP particles. In this system, the biomolecule-stabilized ACP precursors can maintain their amorphous feature for at least 35 days at 37 °C and 120 days at 4 °C (**Figure 6B-6C, Figure S23-S25**). After 42 days of reaction at 37 °C, ACP precursors underwent phase transition and precipitated into HAp crystals, producing needle-like and platelet-like mature morphologies (**Figure 6B-6C**).

**Figure 6:**
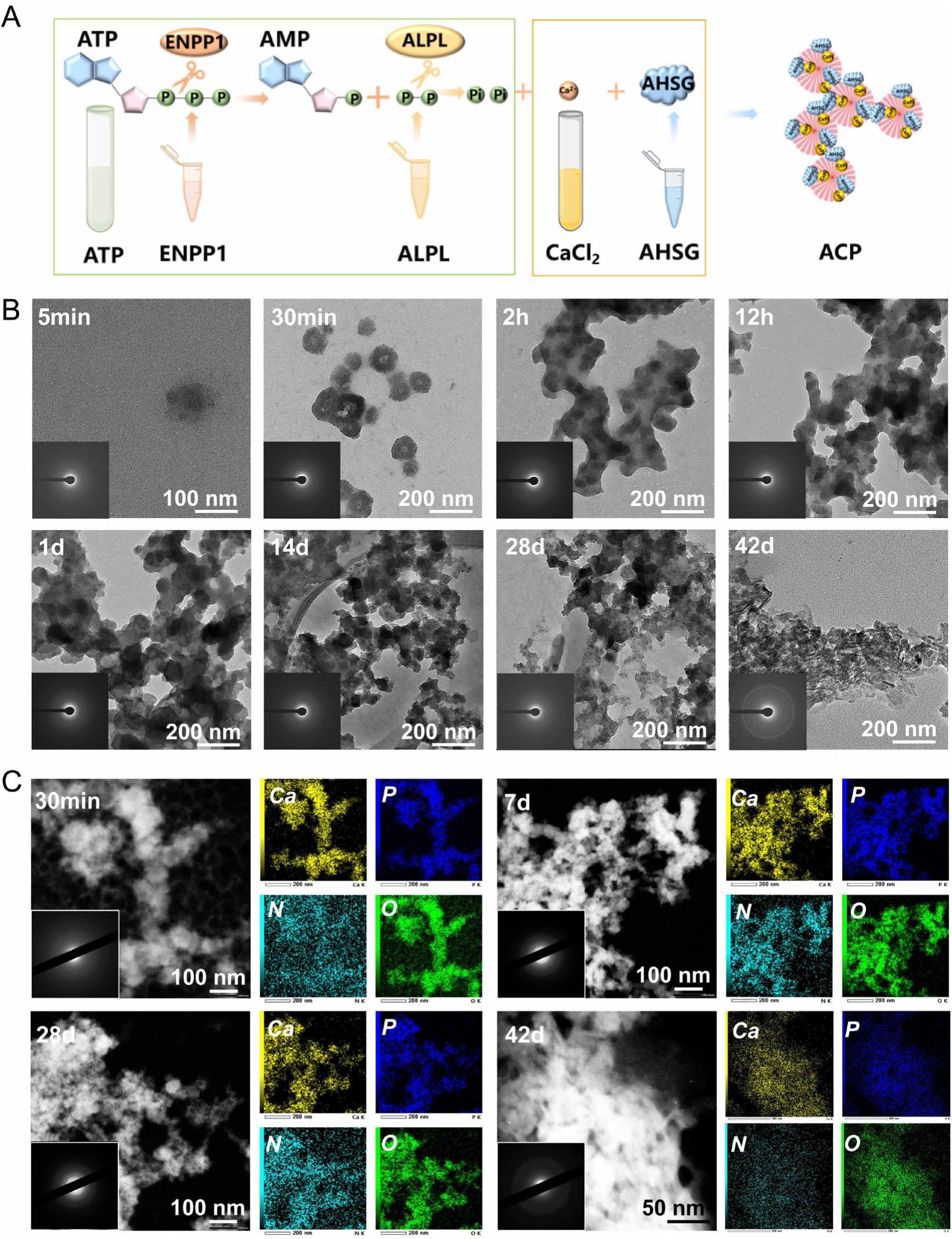
Preparation and characterization of biomolecule-stabilized ACP. A) Schematic showing the preparation of protein-stabilized ACP; B) Representative cryo-TEM and SAED images of prepared ACP after 5min, 2h, 12h, 1d, 5d, 14d, 28d and 42d of reaction at 37 °C, scale bar = 200 nm; C) Representative images of STEM images, EDS mapping images of Ca, P, N, O and SAED pattern of the ACP particles after 30min, 7d, 28d and 42d of reaction at 37 °, scale bar = 100nm/50nm in STEM images, scale bar = 200nm in EDS mapping images.

Although AHSG was suggested to have the ability to form calciprotein particles and restrain CaP precipitation, our results showed that AHSG can only maintain amorphous calciprotein particles less than 15 min (**Figure S26**). The combination of AHSG and ALPL (PPi was provided as phosphorous source) can also stabilize ACP for 3 days at 37 °C. After 5 days, the ACP gradually mineralized into HAp (**Figure S27**). These outcomes collectively reveal, marking a groundbreaking revelation, that hyperstable ACP was successfully fabricated by proteins found at the GP-metaphysis interface in the human knee joint.

## Discussion

Our study revealed the polarized GP mineralization pattern during bone elongation, encompassing the mechanical microenvironment, micro-scale structural and compositional transition, nano-scale crystal assembly, and the underlying regulatory mechanism. Importantly, we highlighted the adaptive nature of bone tissue structure and mineralization in response to external loading, as supported by previous research(*18*, *27*, *30*, *40*). In the case of GPs, the different mechanical properties of the epiphysis and metaphysis contribute to distinct adaptive responses towards mechanical loading, resulting in diverse structural transformations and biomineralization performance at these two soft-hard interfaces, suggesting that applying different patterns of external mechanical loading could be a promising approach for regulating mineralization *in vitro*. Furthermore, the fractal-like hierarchical architecture of minerals at the nano scale in mature human bone tissue has been well investigated(*41*). By demonstrating the microstructure and nano HAp crystal assembly during biomineralization in developing human bone, our research complements the existing knowledge and provides a more comprehensive view of the structural integrity of human bone tissue.

ACP is widely accepted as the vital precursor and intermediate phase during biomineralization. Although the presence of ACP existed in ECM vesicles of developing bone has been reported a few decades ago(*39*), the limited availability of samples and characterization methods have hindered the full investigation of the ACP-like phase in human bone tissue. In this study, we confirmed the existence and transformation of ACP as the mineralization precursor in the ECM during human bone lengthening by multiple high-resolution imaging technologies. It’s widely recognized that ACP precursors are released from intracellular vesicles via exocytosis and organelles such as mitochondria(*42*) and lysosomes(*43*) play a crucial role in this process. One limitation of our research is the lack of investigation into the connection between ACP precursors and cellular behavior during GP mineralization.

Non-collagenous proteins (NCPs) have been regarded as key factors in stabilizing ACP and regulating biomineralization(*44*, *45*). However, the specific proteins involved and their cooperative mechanisms during biomineralization remain unclear. For the first time, we propose a novel concept of “mineralization waves” that govern the growth plate (GP)-guided mineralization process, based on the macromolecule machine regulatory mechanism. Similar to the propagation of sediment waves in a riverbed (**Figure 7**), mineralization waves originate from the GP hypertrophic zone and propagate towards the GP-metaphysis interface. These waves are characterized by the sequential deposition of CaP minerals, with the ACP phase serving as a precursor to form more stable HAp phase. The enzymes at the GP-metaphysis interface creates a dynamic environment that regulates the propagation of these “mineralization waves”. The inhibitory proteins, SPP1 and AHSG, act as “wave attenuators,” slowing down the mineralization process and preventing excessive mineral deposition at the GP-epiphysis interface. When combining with the mineralization-promoting enzymes, ENPP1 and ALPL at the GP-metaphysis interface, these proteins serve as molecular machinery for “wave amplifiers,” accelerating the formation of ACP and its transformation to Hap and facilitating the progression of the mineralization front. Nevertheless, the regulatory mechanism of polarized GP mineralization proposed in our study is speculative, based on the distribution and function of enriched proteins. To further elucidate the *in situ* protein distribution in their biological context and their interaction with resident cells at high resolution, advanced technologies such as correlated light microscopy and electron microscopy (CLEM) need to be employed.

**Figure 7:**
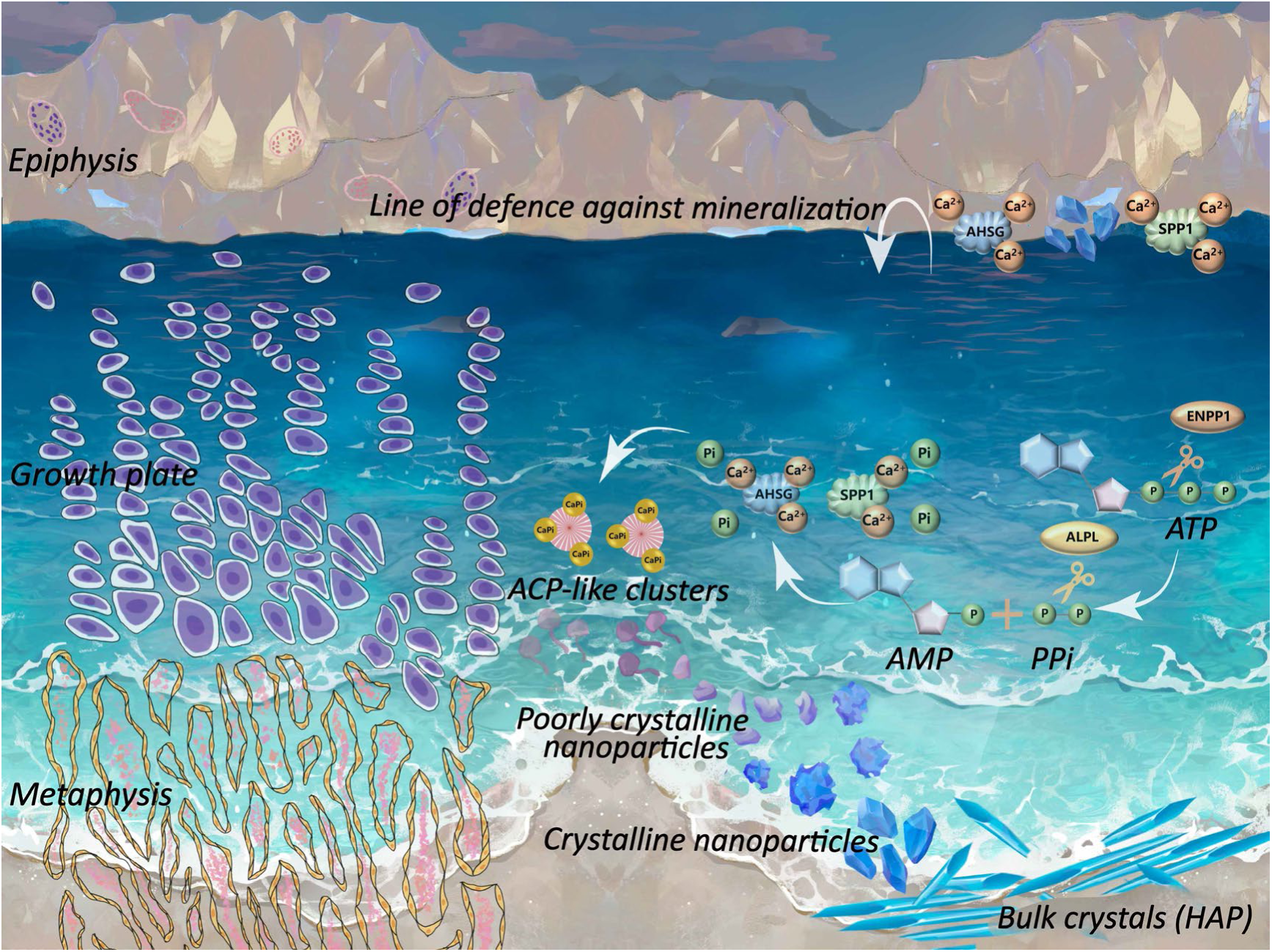
Schematic summary depicting the concept of “mineralization waves” that govern the growth plate (GP)-guided polarized mineralization process. The inhibitory proteins, SPP1 and AHSG, act as “wave attenuators,” forming a line of defence against mineralization in GP-epiphysis interface. When combining with the mineralization-promoting enzymes, ENPP1 and ALPL at the GP-metaphysis interface, these proteins serve as molecular machinery for “wave amplifiers,” accelerating the formation of ACP and its transformation to HAp and facilitating the progression of the mineralization front.

In recent years, constructing ACP *in vitro* to achieve biomimetic mineralization has been a hot topic in the field of bone tissue engineering. ACP has been successfully stabilized by proteins(*7*, *9*), polymers(*45*, *46*), small molecules(*47*) and ions(*5*, *48*). However, the reconstructed ACP stabilized by protein segments and polymers can only maintain its metastable state for several hours or days(*7*, *49*). In addition, the biocompatibility of the ACP is limited by the acidic condition or toxic stabilizing agents(*50*). In this study, by mimicking the cascade processes during GP mineralization, we fabricated ultra-stable ACP that maintained its amorphous state for more than 35 days at 37 °C (more than 120 days at 4 °C) with pH of 7.0-7.4, using a combination of a series of proteins. Stabilizing ACP under mild physiological conditions for such a long period advances our understanding of controlled mineralization processes, preventing pathological mineralization and developing new biomedical materials for in bone regenerative medicine and tissue engineering. By adjusting the concentration and combination of these proteins, it is hoped that ACPs with variable sizes, structures, assemblies and orientations can be fabricated to meet the requirements of different applications in bone regenerative medicine.

## Methods and Materials

### Sample preparation

The growth plate (GP) samples were procured from patients undergoing amputation surgery due to osteosarcoma or trauma (n=6) or from individuals with polydactyly (n=4). Detailed information regarding the samples can be found in **Table S1**. All specimens were carefully selected to ensure the absence of pathological tissue, and a small section was extracted for histological examination to confirm their normalcy. Ethical approval for this study was obtained from the Second Affiliated Hospital of Zhejiang University Ethics Committee (2022LSYD0923) and Second Hospital of Shanxi Medical University Ethic Committee (Ethics No. 2019YX260). The GPs, along with the epiphyseal and metaphyseal tissues, were meticulously harvested, washed with sterile PBS, and stored at -80 °C for subsequent analysis. Each sample was divided into four portions: one for histology and immunofluorescence staining, another for high-resolution analyses (including SEM, Cryo-TEM, FIB, AFM, among others), a third for XRM, and the final portion for LC-MS analysis.

### Histology and immunofluorescence staining of GP, GP-epiphysis interface and GP-metaphysis interface

To prepare for histological and immunofluorescence staining, GP samples underwent fixation in 4% (w/v) paraformaldehyde for 24 to 48 hours. For Safranin O staining (Solarbio), the samples required decalcification in ethylenediaminetetraacetic acid (EDTA) for 3 weeks, followed by dehydration in graded ethanol, clearing in xylene, and paraffin embedding. Samples were sliced (7 μm) using a Leica slicer. For IF staining, non-decalcified samples were cryo-sectioned (10 μm) and blocked in 5% bovine serum albumin (BSA) for 1-2 hours. Then, the samples were left to incubate with primary antibodies overnight at 4°C, followed by PBS washing, and then incubated with fluorescein-conjugated secondary antibodies (Abcam) for 1.5 hours at 37°C. DAPI (Beyotime) was used for visualizing the cell nuclei. Observation was done with a Zeiss LSM 880 confocal microscope. Primary antibodies used were: SPP1 (Santa, sc-21742), AHSG (Proteintech, 66094-1-Ig), ALPL (Proteintech, 11187-1-AP), ENPPI (Abcam, ab223268), PTHrP (Santa, sc-12722), Ki67 (Abcam, ab15580), and COL X (Abcam, ab49945). Mineralized tissue was visualized with 10 μM calcein (DOJINDO) for 20 minutes at room temperature.

### XRM

As described previously, the GP sample (0.5mm long × 0.5mm wide × 1cm height, with GP, epiphysis and metaphysis tissue) was obtained from the normal parts of osteosarcoma patients. The sample was trimmed into a cube with plain epiphysis and metaphysis surfaces. Subsequently, the sample underwent 1% compressive straining in the axial direction and the sample was scanned by an X-ray Microscope (Xradia 620 Versa, Zeiss) before and after exerting the straining, obtaining three-dimensional structural information of the sample. Scan parameter settings are listed as follows: The scan field of view is 11.28 mm high and 11.28 mm wide. Images were collected before and after the GP sample was compressed by 105 μm under a preload of 4.25 N, waiting 15 mins for load relaxation and then micro-CT scanning (5.64 μm per voxel) was performed with the stabilized sample. The total scan time is 2.1 hours. The 3D datasets of static and strained sample were visualized, processed and analyzed using the Digital Volume Correlation (DVC) module in Amira 6.5 (Thermo Fischer). Spatial information of local displacements and strain magnitude were calculated and presented in correlation with the sample morphology according to the software manual.

### AFM

The Cypher atomic force microscope (Oxford instruments Asylum Research, USA) was performed to describe the micromechanical properties of both GP-bone interfaces. The 150 μm cryo-sectioned samples underwent washing and immersion in double distilled water for subsequent measurement. A grid of 32 × 32 pixels in 5 × 5 μm area was measured by silicon nitride cantilevers (AC160TS-R3, Olympus) with a tip radius 9 ± 2 nm and a spring constant 26 N/m. The speed of ramping was set at 3 μm/s until reaching a force of 6 μN, followed by retracting the tip at the same speed. The Hertz model with a conical tip was applied to analyze the Force-Displacement (FD) data for fitting Young’s modulus. The Origin software was used to analyze the data and real map images. 4 areas were measured across each interface.

### SEM, DDC-SEM and EDX analyses

For scanning electron microscopy (SEM) and energy-dispersive X-ray spectroscopy (EDX) analysis, cryo-sectioned GP samples (30 μm) including continuous epiphysis, epiphysis-GP interface, GP, GP-metaphysis interface and metaphysis tissue were rinsed with double distilled water to remove optimal cutting temperature compound (OCT) and then dehydrated by a graded ethanol (20% to 100%) for 30 min in each solution. Next, the samples underwent gold sputtering before observation (HITACHI SU5000), 5 kV accelerating voltage for collecting secondary electrons (SE) and observation of microstructure. Density-dependent color SEM (DDC-SEM) images were acquired at 10 kV, employing both backscattered electron (BSE) mode and SE mode. DDC-SEM images were processed using Image J software, with the green for SE images, the red for BSE images, and two channels stacked to produce single images. Additionally, EDX spectra were collected in point, line, and mapping modes to analyze the elemental compositions of the interfaces. EDX spectra were collected in point, line, and mapping modes to analyze the elemental compositions of the interfaces.

### FIB-SEM

The frozen samples of the GP-metaphysis interface (2mm × 2mm × 2mm) were fixed in a 2.5% (w/v) glutaraldehyde solution for over 12 hours at 4°C. Following this, the samples underwent a series of treatments: rinsing in PBS three times for 15 minutes each, immersion in a solution comprised of 2% osmium tetroxide and 3% potassium ferrocyanide (mixed in a 1:1 ratio) for 1 hour at 4°C, followed by triple rinsing in double distilled water for 10 minutes each. Subsequently, the samples were treated with a 1% (w/v) thiocarbohydrazide solution for 20 minutes, followed by triple rinsing in double distilled water for 10 minutes each. After fixation in a 2% (w/v) osmium tetroxide solution for a duration of 30 minutes at ambient temperature and another round of triple rinsing in double distilled water, the samples were submerged in a 1% (w/v) uranyl acetate solution for an overnight period (over 12 hours) at 4 °C. After washing with double distilled water for 10 minutes, 3 times, the samples underwent dehydration using a series of ethanol concentrations (30%, 50%, 70%, 90%, and 100% twice), with each solution applied for 30 minutes. Then the samples were moved into 100% acetone solution for 20 min, twice. Next, the samples underwent gradient penetration in epoxy resin and embedded in resin.

### FIB-SEM data acquisition, processing, and three-dimensional image reconstruction

The resin blocks were trimmed by ultramicrotome (Leica) until the surface of the sample in the resin blocks became visible. SEM imaging (Thermo Fisher, Teneo VS) was utilized to locate the region of interest, followed by imaging with a dual beam SEM (Thermo Fisher, FIB Helios G3 UC) once the area of interest was identified. After identifying the area of interest, the serial-surface view mode was employed with a slice thickness of 5 nm at 30 keV and 0.79 nA. In each serial face, backscatter mode (BSE) imaging was conducted using a 2 kV acceleration voltage and a current of 0.2 nA, employing an IVD detector. The resolution of each image was 3072 × 2048 pixels, with 15 μs and 4.25 nm per pixel. The image stacks were processed and 3D reconstruction of minerals and collagen fibrils were conducted using Amira 6.5 (Thermo Fisher).

### Stimulated Raman Scattering Microscopy (SRS)

SRS was conducted in liquid state with a commercial SRS microscope (Multimodal Nonlinear Optical Microscopy System, UltraView, Zhendian (Suzhou) Medical Technology Co., Ltd, China), equipped with the InSight X3 (Spectra-Physics/Newport; pulse width, < 120 femtoseconds; tunning range, 680 to 1300 nm) femtosecond laser as light source, supplying tunable pimp beam and fixed Stokes beam. The tested samples were observed by a microscope equipped with 20 X NA 0.8 objective (Olympus) lens and a SRS detection module. The resolution of each image was 512 × 512 pixels. The mapping of ACP (950 cm^-1^), ACP/HAp compounds (955 cm^-1^), HAp (960 cm^-1^), GAG (1410 cm^-1^), protein (1660 cm^-1^ and 2925cm^-1^), lipid (2850 cm^-1^) were analyzed by ImageJ and customized software (SpecFinder). And the final exhibited images were cropped from the original 512 × 512-pixel images.

### Raman spectroscopy

Raman spectroscopy was performed under liquid conditions. The samples were cryo-sectioned into 30 μm without fixation. After washing by double distilled water to remove OCT, the samples underwent observation utilizing a confocal Raman microscope (LabRAM Odyssey) that was outfitted with a 532 nm laser. The spectra were gathered within the range of 200∼1800 cm^-1^ utilizing an electron multiplying charge-coupled device (EMCCD) detector, featuring a spectral resolution of approximately 0.5 cm^-1^. The mapping images were obtained by continuous scanning of 1600 points in 20 μm region each image with an accumulation time of 0.5 s each point. The HAp contents (960 cm^-1^), CO_3_ ^2-^substitution (1071 cm^-1^), mineral crystallinity (full width at half maximum, FWHM of 960 cm^-1^peak), and CO_3_^2-^/PO_4_ ^3-^ratios in mapping images were analyzed by LabSpec software.

### Cryo-TEM observation and tomographic reconstruction

For Cryo-TEM observation and tomographic reconstruction, the GP-Epiphysis interface and GP-metaphysis interface samples (about 2mm each sample, and 3 samples for each interface) including epiphysis, epiphysis-GP interface and GP-metaphysis interface samples including GP, and GP-metaphysis and metaphysis tissue were prepared by high-pressure freezing (HPF) combined with freeze substitution (a mixture of 0.1% osmium tetroxide, 0.1% uranyl acetate, 0.5% glutaraldehyde, 1.5% H_2_O and 100% acetone). Then the specimens were sectioned to 150 nm-thick onto bare 100-mesh copper grids by an ultramicrotome (Leica EM UC7) with cryo-chamber to maintain the samples under -150 °C. The slices were observed in a cryo-TEM (FEI Talos F200C 200kV). For tomographic reconstruction, regions of interest were imaged by tilting the grid in 2° steps from 56° to -56°. The weighted back-projection method was utilized for tomographic reconstruction. Segmentation and 3D visualization were performed using Amira 2019.1 (Visage Imaging Inc., Andover, MA, USA).

### TEM, HR-TEM, SAED, STEM and EELS analyses

For TEM, HR-TEM, STEM, EELS and SAED analyses, the sample preparation is the same as Cryo-TEM. The samples were sectioned to 100 nm-thick onto bare 100-mesh copper grids by ultramicrotome (Leica EM UC7) for further observation. Slices containing GP-Epiphysis and GP-Metaphysis were prepared separately. The slices underwent imaging in a transmission electron microscope (TEM) with spherical aberration correction (FEI Titan G2 80-200), which was furnished with an EELS detector, operating at 80 kV. Regions of interest underwent TEM, HR-TEM, and SAED pattern analyses. The areas for EELS analysis were localized by HAADF-STEM imaging. For EELS mapping, a whole EELS spectrum is acquired at each area. EELS spectra were acquired within the range of 200∼600 eV to investigate the characteristic edges of elements (P, C, Ca, N, O). Each EELS mapping image contains about 20000 spectra. Principal component analysis (PCA) was applied for spectrum calibration, normalization, background subtraction, and processing. Gatan Digital Micrograph software was employed for the analysis of STEM images and EELS.

### Sample preparation for proteomics

Samples (n=3 per group) for proteomics were cryo-sectioned into 100 μm and after sectioning, the slices were segmented by scalpel blade into five parts: epiphysis tissue, epiphysis-GP interface tissue, GP tissue, GP-metaphysis tissue and metaphysis tissue. About six 100 μm-thick slices were required for each group. For digestion, the samples of each group were transferred to 0.6 mL Ep tube and then diluted with 20 μL of 100 mM NH_4_HCO_3_ for 10 min at 95°C. Next, 1 μL of trypsin (1μg/μL) was added to the samples for overnight digestion (12h) at 37°C. Following digestion, any remaining debris was eliminated via centrifugation at 14,000g for 15 minutes at 4°C. The supernatant containing peptides was collected for further experiments. Peptides were quantified using a Nanodrop spectrophotometer (ND-2000C, Thermo) and equal amounts of peptides were taken for desalting. The pH of peptide solution was adjusted to 2-3 by the addition of 20% trifluoroacetic acid (TFA) (Macklin). Desalting was performed using 1.9 μm Reprosil-Pur C18 beads (Dr. Maisch, Ammerbuch, Germany) according to the manufacturer’s instructions, with equilibration by 20 μL of 0.1% TFA. After equilibrating by 20μL 0.1% TFA, the samples were eluted with 0.1% TFA in 80% acetonitrile (Thermo) and subsequently dried using a vacuum concentrator for further analysis.

### LC-MS/MS Analysis

For LC-MS/MS analysis, tryptic peptides were solubilized in 0.1% formic acid (Thermo Fisher) and immediately introduced onto a specialized reversed-phase analytical column filled with 1.9 μm Reprosil-Pur C18 beads (Dr. Maisch, Ammerbuch, Germany). During the process, the gradient elution involved a gradual rise from 3% to 8% solvent (0.1% formic acid in 98% acetonitrile) over 3 minutes, followed by increases to 20% over 37 minutes, then to 30% over 12 minutes, and finally reaching 80% over 4 minutes, maintaining this level for the last 4 minutes. This process was conducted at a consistent flow rate of 300 nL/min using an UltiMate 300 nanoLC system. Next, the peptides underwent NSI source initiation, followed by tandem mass spectrometry analysis using the Orbitrap Exploris 480 (Thermo Fisher), which was integrated with the Ultra Performance Liquid Chromatography (UPLC) system for online coupling. The electrospray voltage was adjusted to 2.0 kV. The full scan mass-to-charge range spanned from 400 to 1200, with intact peptides detected in the Orbitrap at a resolution of 60,000. Peptides were subsequently chosen for LC-MS/MS analysis, employing a normalized collision energy (NCE) setting of 27, and ensuing fragments were identified in the Orbitrap with a resolution of 15,000. A data-dependent approach was employed, alternating between a single MS scan and 20 MS/MS scans with a dynamic exclusion of 30 seconds. Automatic gain control (AGC) was configured at 5E4. The compensation voltages for FAIMS were adjusted to -45V and -65V.

### Database Search for proteomics

The LC-LC-MS/MS data was processed using the MaxQuant search engine (version 1.6.15.0). The tandem mass spectra were compared against the Uniprot Human database concatenated with the reverse decoy database. Trypsin was designated as the cleavage enzyme, permitting a maximum of 2 missed cleavages. The mass deviation for precursor ions was defined as 20 ppm during the initial search and 5 ppm during the primary search, while the mass deviation for fragment ions was set at 0.02 Da. A fixed modification of carbamidomethyl on Cys and a variable modification of oxidation on Met were stipulated. The label-free quantification approach (LFR) was employed, with the Benjamini–Hochberg FDR adjusted to below 1%. Peptides were required to achieve a minimum score exceeding 40.

### Proteomic analysis

The MS/MS data was processed using the MaxQuant search engine (version 1.6.15.0). Initially, the tandem mass spectra were aligned with the Uniprot Human database in conjunction with a reversed decoy database. Trypsin/P was employed as the enzyme for protein cleavage, permitting up to 2 potential missed cleavages. During the initial search, the precursor ion mass tolerance was established to 20 ppm, while for the main search, it was tightened to 5 ppm. Additionally, the tolerance for fragment ion mass was set at 0.02 Da. A fixed modification of carbamidomethyl on cysteine was designated, while oxidation on methionine was treated as a variable modification. The LFQ method was applied for label-free quantification, with the FDR adjusted to < 1%, and a minimum peptide score threshold of > 40 was established. The data obtained were then analyzed utilizing the DEP package within R Studio, with three biological replicates used for analysis. Contaminated samples, reverse data, and duplicated gene names were deleted. Protein rows were filtered to keep only those with at least two out of three valid values observed in individuals per group. Data normalization comprised a variance-stabilizing transformation, succeeded by log2 transformation, while missing values were imputed using the K-nearest neighbors algorithm. Protein expression variances were assessed employing linear models and Empirical Bayes techniques, with significance attributed to fold changes > 2 and adjusted p-values < 0.05.

### TOF-SIMS analysis

The sample preparation for TOF-SIMS mirrors that of FIB-SEM. Utilizing FIB-SEM combined with TOF-SIMS (Thermo Fisher, Helios 5 UX), the distribution and relative abundance of chemical constituents within the samples were analyzed. Throughout the examination, the samples’ surfaces encountered pulses of gallium ion beams. The resulting secondary ions were extracted at a voltage of 10 kV, and a reflection mass spectrometer was utilized to gauge their time of flight from the samples to the detector. Each region measured 100 × 100 μm, comprising 256 × 256 pixels, with 500 scans conducted per area. Both positive and negative ion mass spectra were acquired. The two-dimensional chemical heatmaps with a color-coded scale, showing the intensities of detected secondary ions signal and indicating the relative ion abundance of the scanned area, were analyzed and obtained by TOF-SIMS Explore software.

### Preparation of biomolecule-stabilized ACP

Buffer solution 1 was prepared by dissolving 140mM NaCl (Aladdin) and 50mM Tris in deionized water, and the pH of buffer solution 1 was adjusted to 7.4 with HCl (Diamond). Calcium solution was prepared by dissolving 40 mM CaCl2 (Aladdin) in buffer solution 1. Buffer solution 2 was prepared by dissolving 100mM Tris-HCl (Diamond), and 5mM MgCl2 (Sigma-Aldrich) in deionized water, and the pH of buffer solution 2 was adjusted to 9.0 with NaOH (Sigma-Aldrich). Thereafter, all approaches were conducted in a clean bench. Buffer solution 1, calcium solution, and buffer solution 2 were sterilized by percolating the solutions through a 0.22µm membrane filter. AHSG solution as well as ALPL solution was prepared by dissolving 2mg/mL active recombinant human fetuin A/AHSG protein, and 1mg/mL recombinant human alkaline phosphatase protein in buffer solution 1, respectively. ENPP1 solution was prepared by dissolving 0.5mg/mL recombinant human ENPP1 protein in buffer solution 2. AHSG, ALPL and ENPP1 were purchased from ABclonal.

In reaction solution A, 5µL 100mM ATP solution (Novoprotein), 32uL ENPP1 solution, and 3uL buffer solution 2 were mixed and incubated at 37°C for 1h. Then 10uL ALPL solution was added, followed by a 10-minute incubation. In reaction solution B, 5µL AHSG solution, 25µL calcium solution, and 20µL buffer solution 1 were mixed and incubated at 37°C for 10 minutes. The 50µL reaction solution A and reaction solution B were mixed in a 1:1 ratio and incubated at 37°C. At 5min, 15min, 30min, 1h, 2h, 12h, 1d, 3d, 5d, 7d, 10d, 14d, 21d, 28d, 35d and 42d, samples of the mixture were collected with a 300-mesh gold support grid (Zhongjingkeyi Technology Co., China) for TEM.

### Cryo-TEM, STEM, SAED and EDS mapping of ACP

The samples of reconstructed ACP for Cryo-TEM, STEM imaging and EDS mapping were prepared as said above. The ACP was collected in 300-mesh gold support grids for further cryo-TEM and STEM observation. The ACP samples were observed in a cryo-TEM (FEI Talos F200C) at 200 kV. The samples for STEM were imaged in a field emission TEM (JEOL JEM-F200) at 80 kV. STEM-EDX mapping (Ca, P, N, C) and SAED were performed in the areas of interest.

## Data availability

The LC-MS data generated in this study are available upon reasonable request to the corresponding author.

## Acknowledgements

The authors acknowledged the financial support from the National Key Research and Development Program of China (2023YFB3813000), and the National Natural Sciences Foundation of China (No. T2121004, 82394441, 92268203, 32371411), and the Key Research and Development Program of Zhejiang (2024SSYS0026). The authors would extend their gratitude to Mr. Jiadan Wu and Ms. Junyan Xie (The Second Affiliated Hospital, Zhejiang University) for their assistance on growth plate samples collection. The authors would like to thank Mr. Lu Lan and Mr. Shoupu Yi (Multimodal Nonlinear Optical Microscopy System, UltraView, Zhendian, Suzhou) for their assistance on stimulated Raman scattering microscopy. The authors also thank Mr. Jiansheng Guo (Center of Cryo-Electron Microscopy, Zhejiang University) for his assistance with FIB-SEM, Ms. Lingyun Wu (Center of Cryo-Electron Microscopy, Zhejiang University) for her assistance with Cryo-TEM and Beibei Wang for her assistance with TEM and ultrathin slicing. The authors would like to thank Ms. Guoqing Zhu from the Center of Electron Microscopy of Zhejiang University for her technical assistance on spherical aberration corrected TEM (FEI Titan G2 80-200) characterization and Ms. Qingyun Lin from the Center of Electron Microscopy of Zhejiang University for her technical assistance on F20 TEM characterization. The authors also thank Mr. Pengda Zou and Ms. Minghui Li (Mass Spectrometry Core Facilities, The First Affiliated Hospital, Zhejiang University School of Medicine) for their assistance with LC-MS. The authors also thank Mrs. Chunjie Cao, Ms. Biyu Chen and Ms. Xi Lin from Carl Zeiss AG for their assistance in XRM and data analysis.

## Author contributions

Chang Xie, Wenyue Li contributed equally to this work.

Conceptualization: Chang Xie, Xiaozhao Wang, Hongwei Ouyang

Methodology: Chang Xie, Wenyue Li, Boxuan Wu, Hongxu Meng, Yiyang Yan

Investigation: Chang Xie, Wenyue Li, Xudong Yao, Boxuan Wu

Resources: Wangping Duan, Yan Wu

Visualization: Chang Xie, Wenyue Li, Xudong Yao, Boxuan Wu, Renwei Mao, Yiyang Yan

Supervision: Hongwei Ouyang, Xiaozhao Wang

Writing: Chang Xie, Wenyue Li, Xudong Yao, Boxuan Wu, Xianzhu Zhang, Xiaozhao Wang, Hongwei Ouyang

## References

1. A. Arnold, E. Dennison, C. S. Kovacs, M. Mannstadt, R. Rizzoli, M. L. Brandi, B. Clarke, R. V. Thakker, Hormonal regulation of biomineralization. Nat. Rev. Endocrinol. 17, 261–275 (2021).

2. A. Gal, R. Wirth, J. Kopka, P. Fratzl, D. Faivre, A. Scheffel, Macromolecular recognition directs calcium ions to coccolith mineralization sites. Science (80-. ). 353, 590–593 (2016).

3. A. Dey, P. H. H. Bomans, F. A. Müller, J. Will, P. M. Frederik, G. De With, N. A. J. M. Sommerdijk, The role of prenucleation clusters in surface-induced calcium phosphate crystallization. Nat. Mater. 9, 1010–1014 (2010).

4. A. Lotsari, A. K. Rajasekharan, M. Halvarsson, M. Andersson, Transformation of amorphous calcium phosphate to bone-like apatite. Nat. Commun. 9 (2018), doi:10.1038/s41467-018-06570-x.

5. S. Chen, D. Liu, L. Fu, B. Ni, Z. Chen, J. Knaus, E. V Sturm, B. Wang, H. J. Haugen, H. Yan, H. Cölfen, B. Li, Formation of Amorphous Iron-Calcium Phosphate with High Stability. 2301422, 1–11 (2023).

6. N. Golafshan, E. Vorndran, S. Zaharievski, H. Brommer, F. B. Kadumudi, A. Dolatshahi-Pirouz, U. Gbureck, R. van Weeren, M. Castilho, J. Malda, Tough magnesium phosphate-based 3D-printed implants induce bone regeneration in an equine defect model. Biomaterials. 261 (2020), doi:10.1016/j.biomaterials.2020.120302.

7. R. Chang, Y. J. Liu, Y. L. Zhang, S. Y. Zhang, B. B. Han, F. Chen, Y. X. Chen, Phosphorylated and Phosphonated Low-Complexity Protein Segments for Biomimetic Mineralization and Repair of Tooth Enamel. Adv. Sci. 9, 1–16 (2022).

8. D. Wang, J. Deng, X. Deng, C. Fang, X. Zhang, P. Yang, Controlling Enamel Remineralization by Amyloid-Like Amelogenin Mimics. Adv. Mater. 32, 1–13 (2020).

9. W. Jahnen-Dechent, A. Pasch, Solving the insoluble: calciprotein particles mediate bulk mineral transport. Kidney Int. 103, 663–665 (2023).

10. M. juan Shen, K. Jiao, C. yu Wang, H. Ehrlich, M. chen Wan, D. xiao Hao, J. Li, Q. qian Wan, L. Tonggu, J. fei Yan, K. yan Wang, Y. xuan Ma, J. hua Chen, F. R. Tay, L. na Niu, Extracellular DNA: A Missing Link in the Pathogenesis of Ectopic Mineralization. Adv. Sci. 9, 1–13 (2022).

11. M. juan Shen, C. yu Wang, D. xiao Hao, J. xin Hao, Y. fei Zhu, X. xiao Han, L. Tonggu, J. hua Chen, K. Jiao, F. R. Tay, L. na Niu, Multifunctional Nanomachinery for Enhancement of Bone Healing. Adv. Mater. 34, 1–12 (2022).

12. A. Behrooz, P. Kask, J. Meganck, J. Kempner, Automated quantitative bone analysis in in vivo x-ray micro-computed tomography. IEEE Trans. Med. Imaging. 36, 1955–1965 (2017).

13. Y. Ağirdil, The growth plate: A physiologic overview. EFORT Open Rev. 5, 498–507 (2020).

14. A. S. Tiffany, B. A. C. Harley, Growing Pains: The Need for Engineered Platforms to Study Growth Plate Biology. Adv. Healthc. Mater. 11, 1–19 (2022).

15. T. Voller, P. Cameron, J. Watson, J. Phadnis, The growth plate: anatomy and disorders. Orthop. Trauma. 34, 135–140 (2020).

16. K. Mizuhashi, W. Ono, Y. Matsushita, N. Sakagami, A. Takahashi, T. L. Saunders, T. Nagasawa, H. M. Kronenberg, N. Ono, Resting zone of the growth plate houses a unique class of skeletal stem cells. Nature. 563, 254–258 (2018).

17. P. T. Newton, L. Li, B. Zhou, C. Schweingruber, M. Hovorakova, M. Xie, X. Sun, L. Sandhow, A. V. Artemov, E. Ivashkin, S. Suter, V. Dyachuk, M. El Shahawy, A. Gritli-Linde, T. Bouderlique, J. Petersen, A. Mollbrink, J. Lundeberg, G. Enikolopov, H. Qian, K. Fried, M. Kasper, E. Hedlund, I. Adameyko, L. Sävendahl, A. S. Chagin, A radical switch in clonality reveals a stem cell niche in the epiphyseal growth plate. Nature. 567, 234–238 (2019).

18. P. Christen, K. Ito, R. Ellouz, S. Boutroy, E. Sornay-Rendu, R. D. Chapurlat, B. Van Rietbergen, Bone remodelling in humans is load-driven but not lazy. Nat. Commun. 5 (2014), doi:10.1038/ncomms5855.

19. W. J. E. M. Habraken, J. Tao, L. J. Brylka, H. Friedrich, L. Bertinetti, A. S. Schenk, A. Verch, V. Dmitrovic, P. H. H. Bomans, P. M. Frederik, J. Laven, P. Van Der Schoot, B. Aichmayer, G. De With, J. J. DeYoreo, N. A. J. M. Sommerdijk, Ion-association complexes unite classical and non-classical theories for the biomimetic nucleation of calcium phosphate. Nat. Commun. 4 (2013), doi:10.1038/ncomms2490.

20. J. J. De Yoreo, P. U. P. A. Gilbert, N. A. J. M. Sommerdijk, R. L. Penn, S. Whitelam, D. Joester, H. Zhang, J. D. Rimer, A. Navrotsky, J. F. Banfield, A. F. Wallace, F. M. Michel, F. C. Meldrum, H. Cölfen, P. M. Dove, Crystallization by particle attachment in synthetic, biogenic, and geologic environments. Science (80-. ). 349 (2015), doi:10.1126/science.aaa6760.

21. H. Fleisch, R. G. G. Russell, F. Straumann, Effect of pyrophosphate on hydroxyapatite and its implications in calcium homeostasis. Nature. 212, 901– 903 (1966).

22. E. J. Novais, R. Narayanan, J. A. Canseco, K. van de Wetering, C. K. Kepler, A. S. Hilibrand, A. R. Vaccaro, M. V. Risbud, A new perspective on intervertebral disc calcification—from bench to bedside. Bone Res. 12, 1–12 (2024).

23. C. Goettsch, A. Strzelecka-Kiliszek, L. Bessueille, T. Quillard, L. Mechtouff, S. Pikula, E. Canet-Soulas, M. J. Luis, C. Fonta, D. Magne, TNAP as a therapeutic target for cardiovascular calcification: A discussion of its pleiotropic functions in the body. Cardiovasc. Res. 118, 84–96 (2022).

24. N. Reznikov, J. A. M. Steele, P. Fratzl, M. M. Stevens, A materials science vision of extracellular matrix mineralization. Nat. Rev. Mater. 1 (2016), doi:10.1038/natrevmats.2016.41.

25. F. Nudelman, K. Pieterse, A. George, P. H. H. Bomans, H. Friedrich, L. J. Brylka, P. A. J. Hilbers, G. De With, N. A. J. M. Sommerdijk, The role of collagen in bone apatite formation in the presence of hydroxyapatite nucleation inhibitors. Nat. Mater. 9, 1004–1009 (2010).

26. K. Madi, K. A. Staines, B. K. Bay, B. Javaheri, H. Geng, A. J. Bodey, S. Cartmell, A. A. Pitsillides, P. D. Lee, In situ characterization of nanoscale strains in loaded whole joints via synchrotron X-ray tomography. *Nat*. Biomed. Eng. 4, 343–354 (2020).

27. W. Huang, M. Shishehbor, N. Guarín-Zapata, N. D. Kirchhofer, J. Li, L. Cruz, T. Wang, S. Bhowmick, D. Stauffer, P. Manimunda, K. N. Bozhilov, R. Caldwell, P. Zavattieri, D. Kisailus, A natural impact-resistant bicontinuous composite nanoparticle coating. Nat. Mater. 19, 1236–1243 (2020).

28. S. Kim, A. U. Regitsky, J. Song, J. Ilavsky, G. H. McKinley, N. Holten-Andersen, In situ mechanical reinforcement of polymer hydrogels via metal-coordinated crosslink mineralization. Nat. Commun. 12, 1–10 (2021).

29. M. Milazzo, G. S. Jung, S. Danti, M. J. Buehler, Mechanics of Mineralized Collagen Fibrils upon Transient Loads. ACS Nano. 14, 8307–8316 (2020).

30. J. H. Chen, C. Liu, L. You, C. A. Simmons, Boning up on Wolff’s Law: Mechanical regulation of the cells that make and maintain bone. J. Biomech. 43, 108–118 (2010).

31. K. Nitiputri, Q. M. Ramasse, H. Autefage, C. M. McGilvery, S. Boonrungsiman, N. D. Evans, M. M. Stevens, A. E. Porter, Nanoanalytical Electron Microscopy Reveals a Sequential Mineralization Process Involving Carbonate-Containing Amorphous Precursors. ACS Nano. 10, 6826–6835 (2016).

32. A. Akiva, M. Kerschnitzki, I. Pinkas, W. Wagermaier, K. Yaniv, P. Fratzl, L. Addadi, S. Weiner, Mineral Formation in the Larval Zebrafish Tail Bone Occurs via an Acidic Disordered Calcium Phosphate Phase. J. Am. Chem. Soc. 138, 14481–14487 (2016).

33. J. A. Stammeier, B. Purgstaller, D. Hippler, V. Mavromatis, M. Dietzel, In-situ Raman spectroscopy of amorphous calcium phosphate to crystalline hydroxyapatite transformation. MethodsX. 5, 1241–1250 (2018).

34. A. Akiva, G. Malkinson, A. Masic, M. Kerschnitzki, M. Bennet, P. Fratzl, L. Addadi, S. Weiner, K. Yaniv, On the pathway of mineral deposition in larval zebrafish caudal fin bone. Bone. 75, 192–200 (2015).

35. R. L. Penn, J. F. Banfield, Imperfect oriented attachment: Dislocation generation in defect-free nanocrystals. Science (80-. ). 281, 969–971 (1998).

36. W. Jahnen-Dechent, E. R. Smith, Nature’s remedy to phosphate woes: calciprotein particles regulate systemic mineral metabolism. Kidney Int. 97, 648–651 (2020).

37. K. Kato, H. Nishimasu, S. Okudaira, E. Mihara, R. Ishitani, J. Takagi, J. Aoki, O. Nureki, Crystal structure of Enpp1, an extracellular glycoprotein involved in bone mineralization and insulin signaling. Proc. Natl. Acad. Sci. U. S. A. 109, 16876–16881 (2012).

38. C. Bussola Tovani, A. Gloter, T. Azaïs, M. Selmane, A. P. Ramos, N. Nassif, Formation of stable strontium-rich amorphous calcium phosphate: Possible effects on bone mineral. Acta Biomater. 92, 315–324 (2019).

39. S. Chen, D. Liu, L. Fu, B. Ni, Z. Chen, J. Knaus, E. V. Sturm, B. Wang, H. J. Haugen, H. Yan, H. Cölfen, B. Li, Formation of Amorphous Iron-Calcium Phosphate with High Stability. Adv. Mater. 35, 1–11 (2023).

40. M. Cusack, A. Freer, Biomineralization: Elemental and organic influence in carbonate systems. Chem. Rev. 108, 4433–4454 (2008).

41. N. Reznikov, M. Bilton, L. Lari, M. M. Stevens, R. Kröger, Fractal-like hierarchical organization of bone begins at the nanoscale. Science (80-. ). 360 (2018), doi:10.1126/science.aao2189.

42. D. Pei, J. Sun, C. Zhu, F. Tian, K. Jiao, M. R. Anderson, C. Yiu, C. Huang, C. Jin, B. E. Bergeron, J. Chen, F. R. Tay, L. Niu, Contribution of Mitophagy to Cell-Mediated Mineralization : Revisiting a 50-Year-Old Conundrum. 1800873 (2018), doi:10.1002/advs.201800873.

43. T. Iwayama, T. Okada, T. Ueda, K. Tomita, S. Matsumoto, M. Takedachi, S. Wakisaka, T. Noda, T. Ogura, T. Okano, P. Fratzl, T. Ogura, S. Murakami, Osteoblastic lysosome plays a central role in mineralization. 10, 1–9 (2019).

44. G. Thrivikraman, A. Athirasala, R. Gordon, L. Zhang, R. Bergan, D. R. Keene, J. M. Jones, H. Xie, Z. Chen, J. Tao, B. Wingender, L. Gower, J. L. Ferracane, L. E. Bertassoni, Rapid fabrication of vascularized and innervated cell-laden bone models with biomimetic intrafibrillar collagen mineralization. Nat. Commun. 10 (2019), doi:10.1038/s41467-019-11455-8.

45. H. Wu, C. Shao, J. Shi, Z. Hu, Y. Zhou, Z. Chen, R. Tang, Z. Xie, W. Jin, Hyaluronic acid-mediated collagen intrafibrillar mineralization and enhancement of dentin remineralization. Carbohydr. Polym. 319, 121174 (2023).

46. J. xin Hao, Q. qian Wan, Z. Mu, J. ting Gu, W. wei Yu, W. Qin, Y. tao Li, C. yu Wang, Y. xuan Ma, K. Jiao, F. Tay, L. Niu, A seminal perspective on the role of chondroitin sulfate in biomineralization. Carbohydr. Polym. 310, 120738 (2023).

47. C. Shao, R. Zhao, S. Jiang, S. Yao, Z. Wu, B. Jin, Y. Yang, H. Pan, R. Tang, Citrate Improves Collagen Mineralization via Interface Wetting: A Physicochemical Understanding of Biomineralization Control. Adv. Mater. 30, 1–7 (2018).

48. Y. Li, Y. Kong, B. Xue, J. Dai, G. Sha, H. Ping, L. Lei, W. Wang, K. Wang, Z. Fu, Mechanically Reinforced Artificial Enamel by Mg2+-Induced Amorphous Intergranular Phases. ACS Nano. 16, 10422–10430 (2022).

49. J. He, J. Yang, M. Li, Y. Li, Y. Pang, J. Deng, X. Zhang, W. Liu, Polyzwitterion Manipulates Remineralization and Antibiofilm Functions against Dental Demineralization. ACS Nano. 16, 3119–3134 (2022).

50. S. Yao, X. Lin, Y. Xu, Y. Chen, P. Qiu, C. Shao, B. Jin, Z. Mu, N. A. J. M. Sommerdijk, R. Tang, Osteoporotic Bone Recovery by a Highly Bone-Inductive Calcium Phosphate Polymer-Induced Liquid-Precursor. Adv. Sci. 6 (2019), doi:10.1002/advs.201900683.

