## Supplemental figures and tables for "Bio-macromolecular assembly machine for human limb lengthening"

### Supplementary Figures:

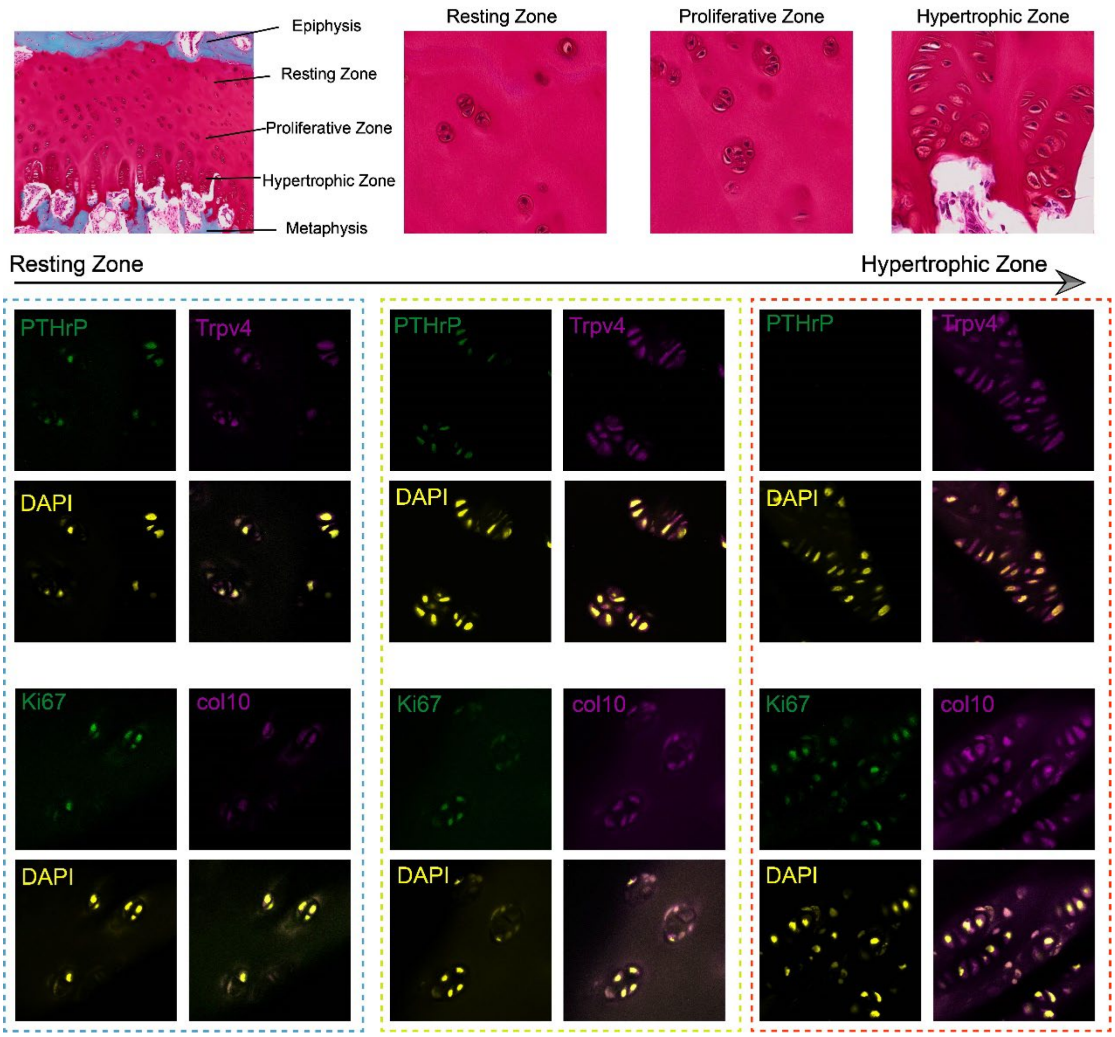


Figure S1: The immunofluorescent staining of PTHrP, Trpv4, Ki67 and col 10 from resting zone to hypertrophic zone, showing the highly ordered structure and different physiological states of zones within GP.

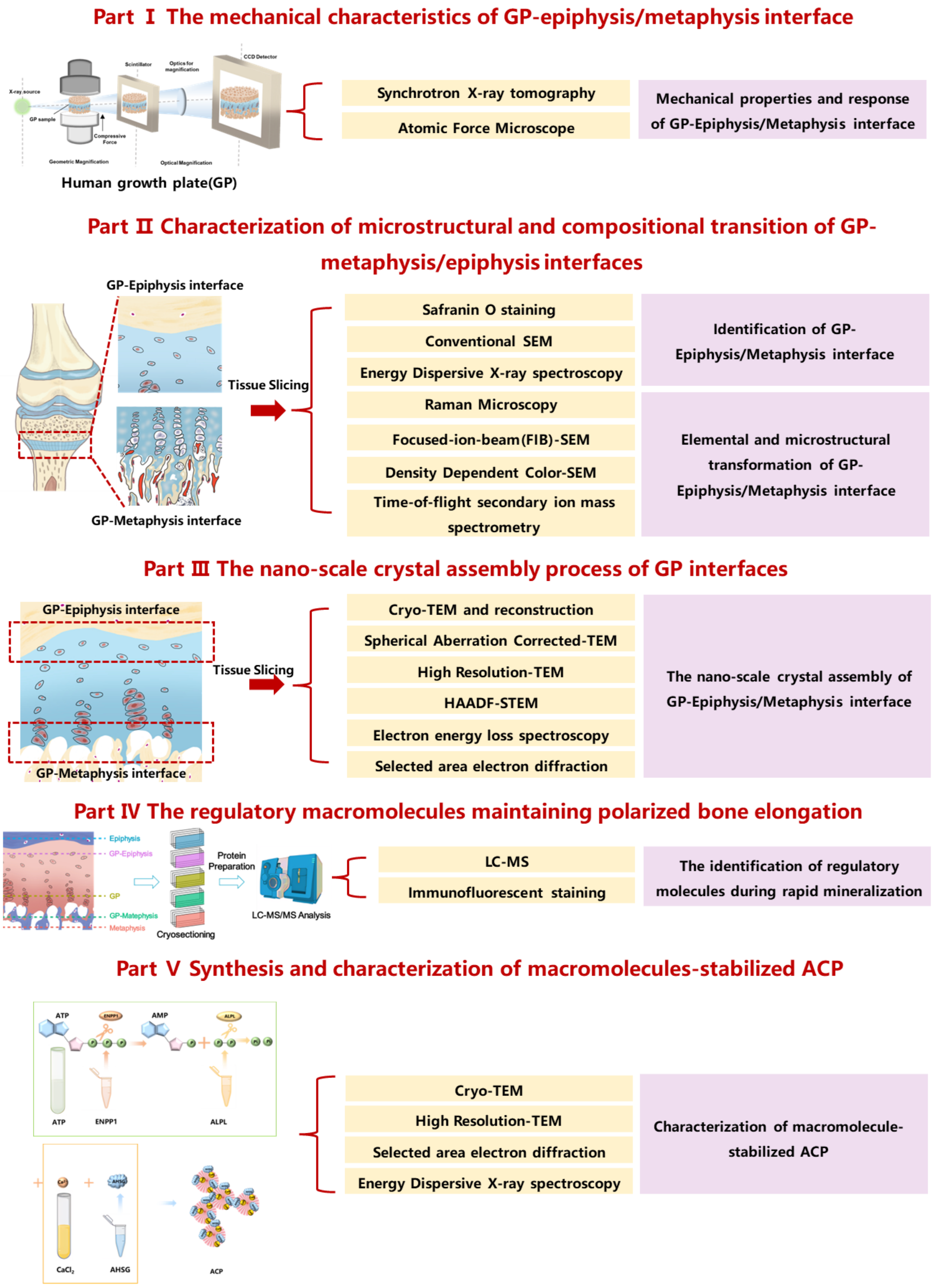


Figure S2: The workflow of the research.

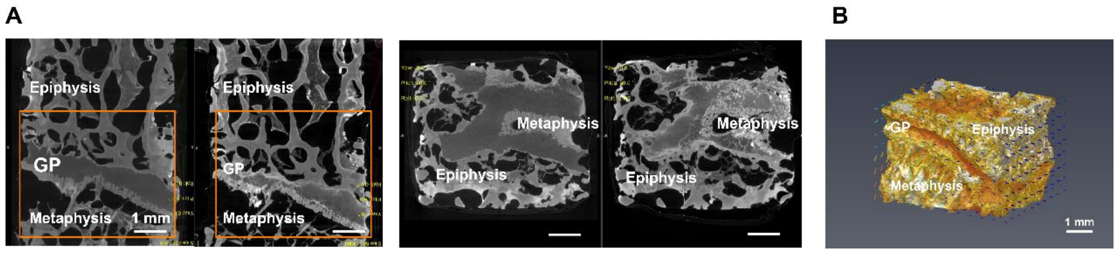

Figure S3: A) Selected CT images of GP tissue before and after compression via synchrotron XRM, scale bar = 1mm; B) The reconstructed CT image of GP after compression, the arrows represent the displacement direction, scale bar = 1mm.

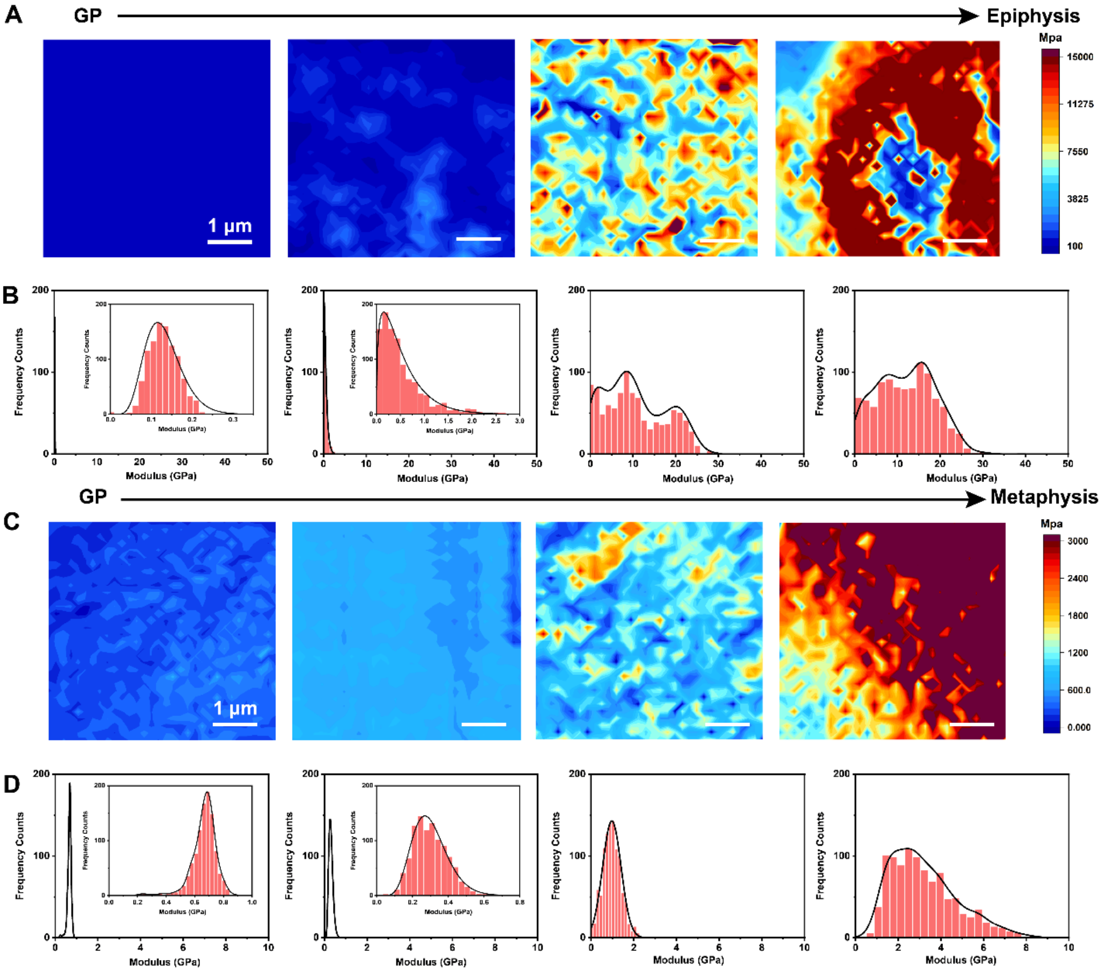


Figure S4: A-B) AFM stiffness maps and corresponding stiffness distribution of selected regions from GP to epiphysis, scale bar = 1μm; C-D) AFM stiffness maps and corresponding stiffness distribution of selected regions from GP to metaphysis, scale bar = 1μm.

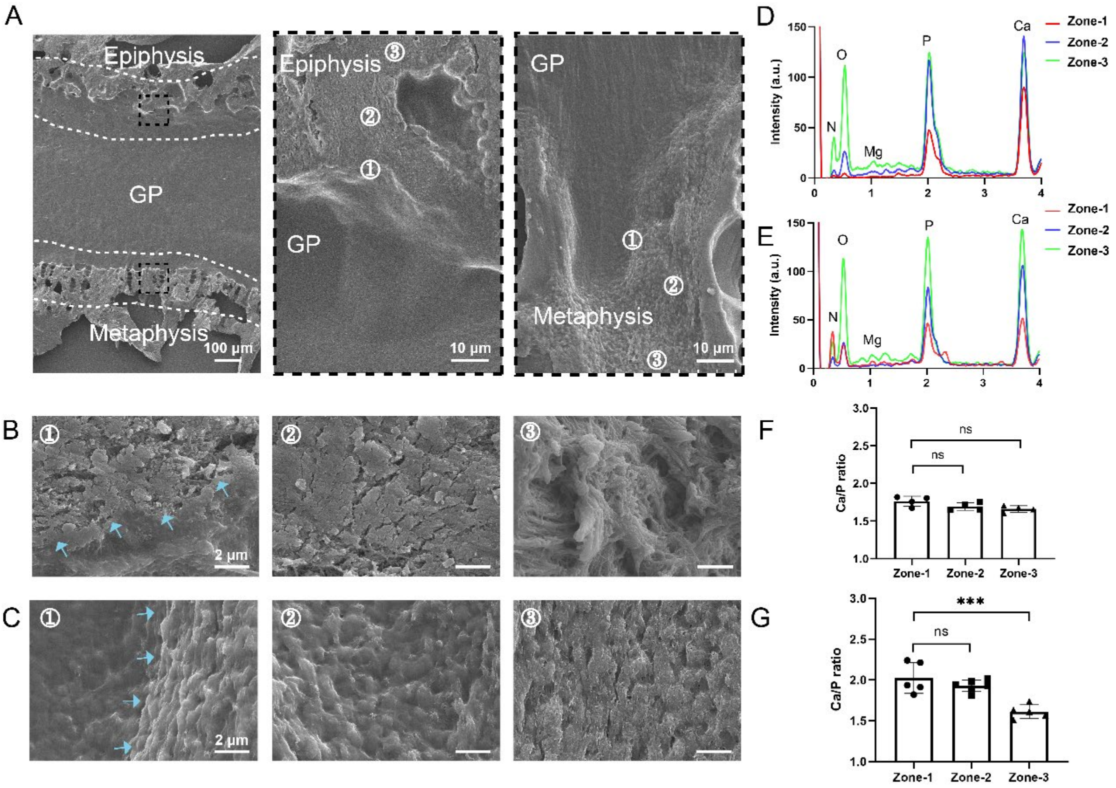


Figure S5: A) Representative SEM images of GP and enlarged images of GP-epiphysis interface and GP-epiphysis interface, scale bar = 100 μm; B) The enlarged SEM images of zone 1-3 in GP-epiphysis interface; C) The enlarged SEM images of zone 1-3 in GP-metaphysis interface, blue arrows pointing to the soft-hard interface in B-C), scale bar = 2 μm; C-D) Chemical composition of C) GP-epiphysis and D) GP-metaphysis interface in zone 1-3; E-F) The corresponding Ca/P ratios of minerals within each zone from GP to E) epiphysis and F) metaphysis.

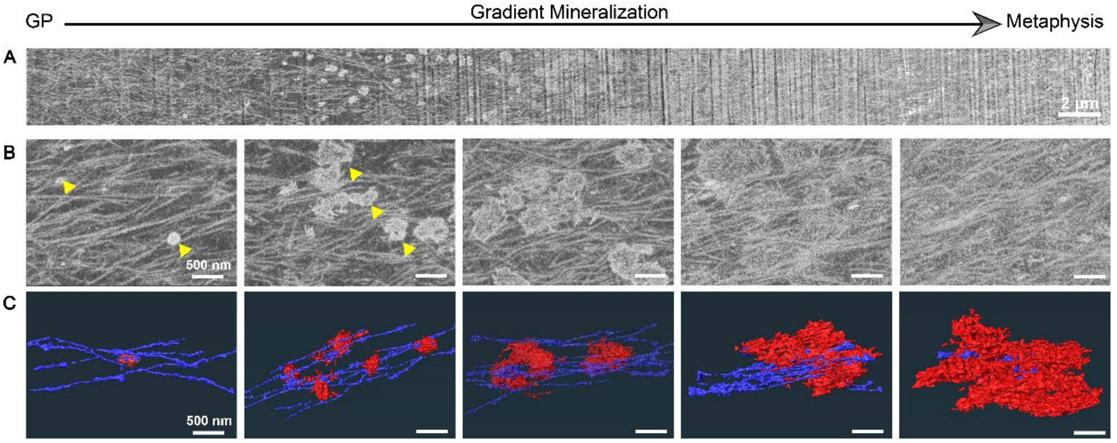


Figure S6: A-B) FIB-SEM (backscatter electron mode) images of GP-metaphysis interface from GP cartilage to mineralized metaphysis region, scale bar = 2 μm in A) and 500 nm in B), triangles pointing minerals; 3D reconstruction images showing the collagen fibers (violet) and minerals (red) across GP-metaphysis interface, scale bar = 500 nm.

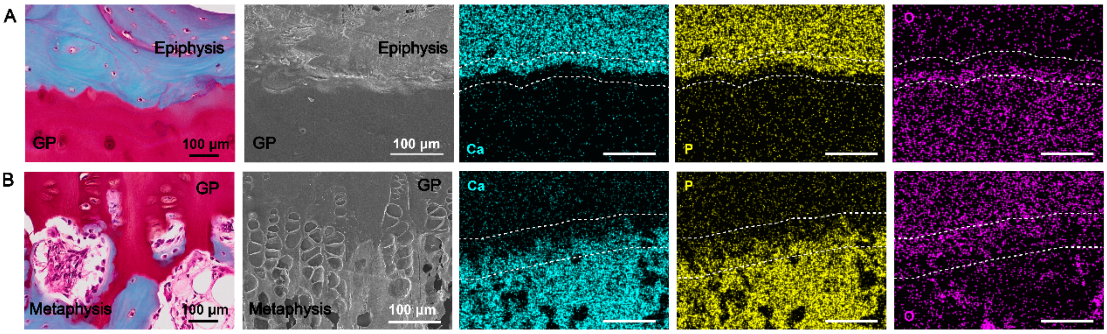


Figure S7: A-B) Enlarged SO staining image and SEM image of A) the GP-epiphysis interface and B) the GP-metaphysis interface, as well as corresponding EDX mapping of Ca, P and O, the dotted lines showing the regions of elemental transition, scale bar = 100µm; SO, Safranin-O staining.

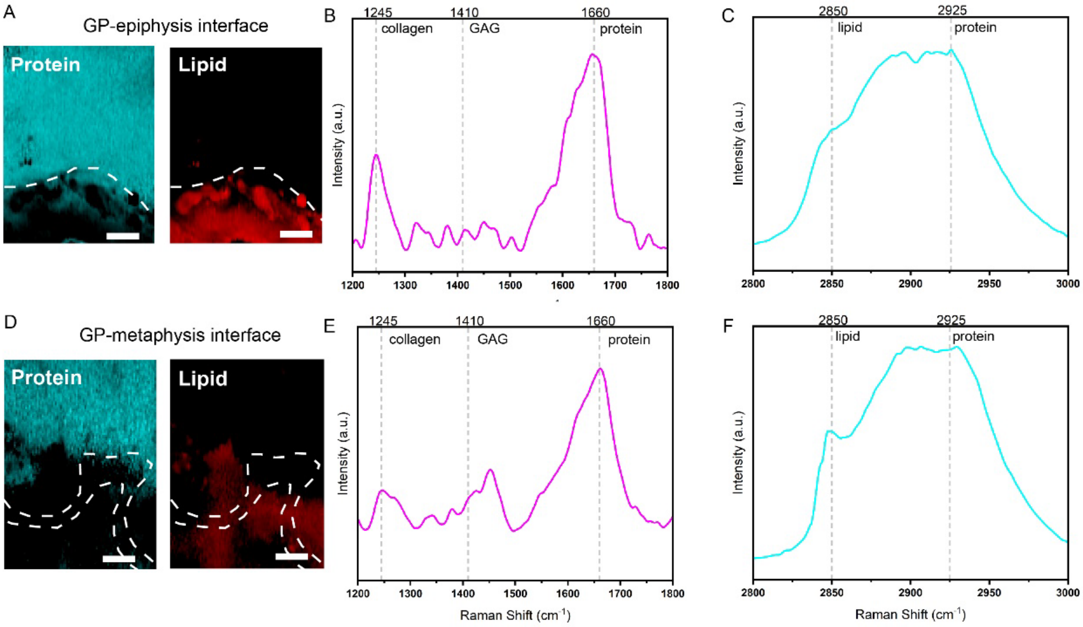


Figure S8: SRS imaging of GP-epiphysis interface A) and GP-metaphysis interface D) showing the band intensity associated with protein (1660 cm$^{-1}$) and lipid (2850 cm$^{-1}$), scale bar = 10 µm; Representative Raman spectra of GP-epiphysis interface B, C) and GP-metaphysis interface E, F) from SRS imaging with mark signatures for collagen (1245 cm$^{-1}$), GAG (1410 cm$^{-1}$), protein (1660 cm$^{-1}$ and 2925 cm$^{-1}$) and lipid (2850 cm$^{-1}$).

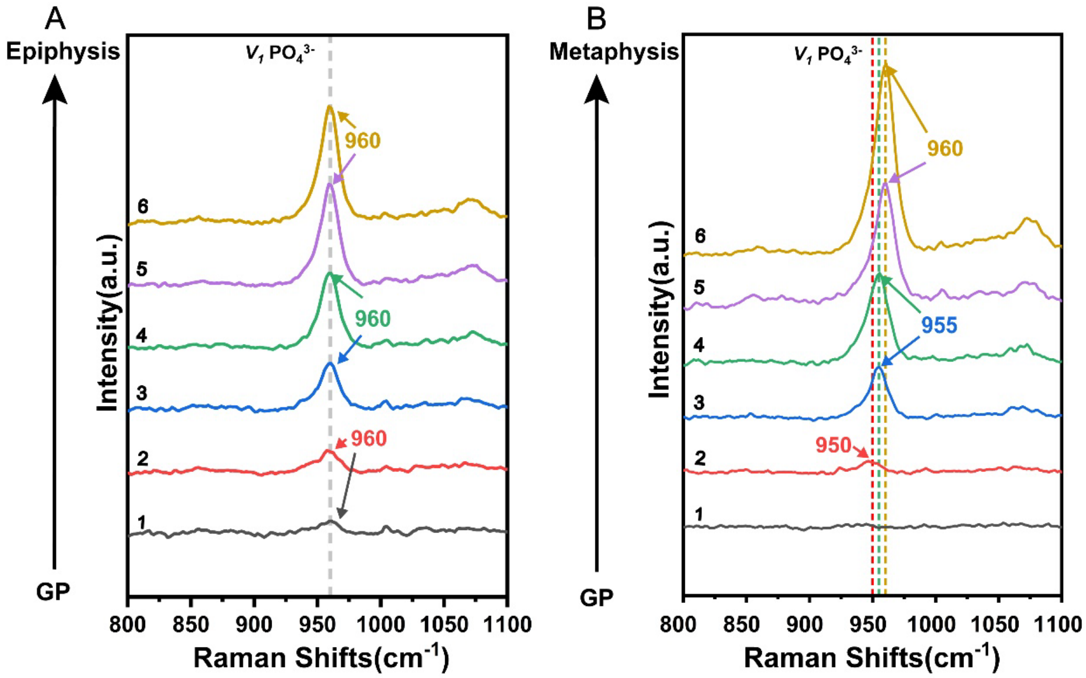


Figure S9: Enlarged images of Raman spectra collected in 800 ~1100 cm$^{-1}$ from GP to A) epiphysis/ B) metaphysis at different area (zone 1-6) in Figure 3A and 3B.

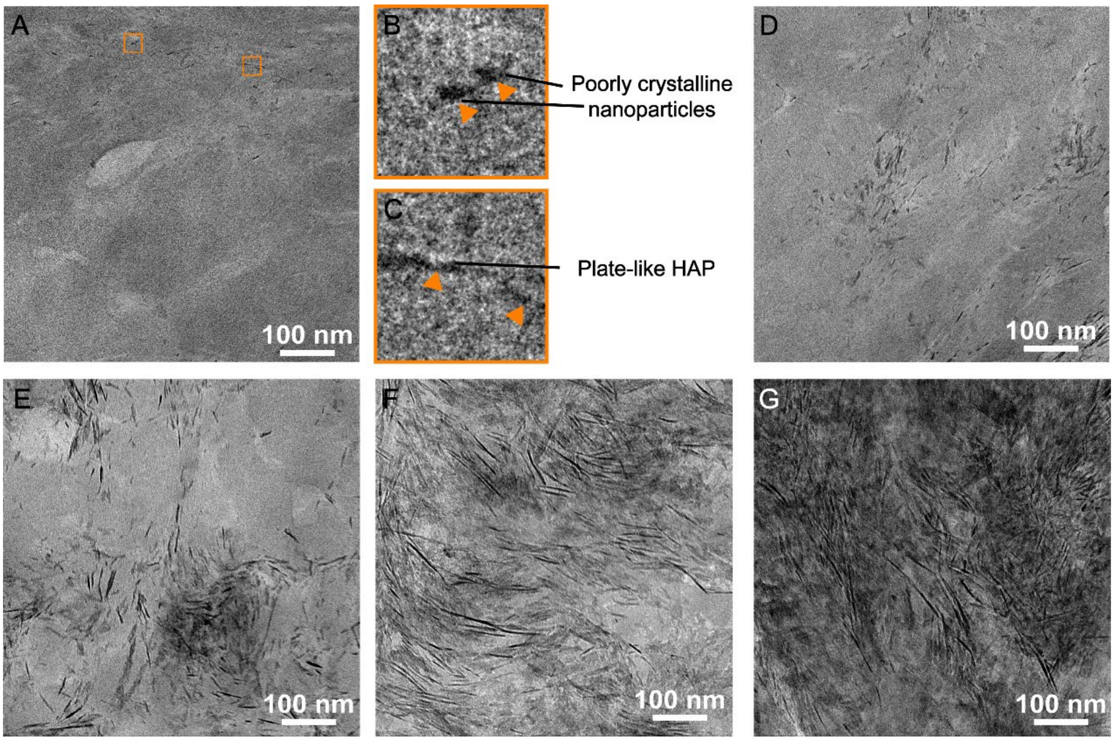


Figure S10: Representative cryo-TEM images in GP-epiphysis interface of tibia sample, from poorly crystalline HAp nanoparticles and plate-like HAp to fully mineralized epiphysis region, scale bar = 100 nm.

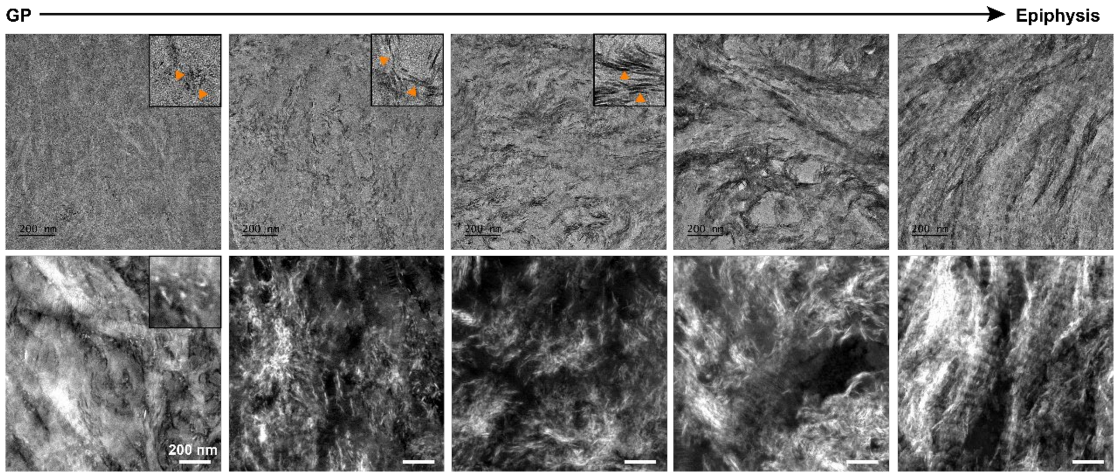


Figure S11: Representative TEM and STEM images of tibia GP sample showing the mineralization process of GP-epiphysis interface from poorly crystalline nanoparticles to fully mineralized metaphysis tissue, scale bar = 200 nm.

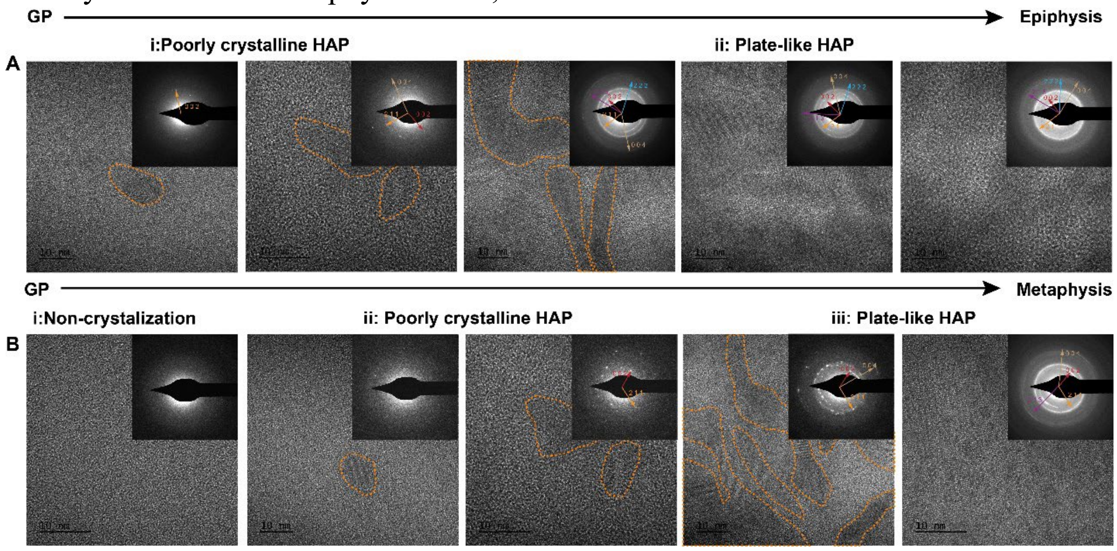


Figure S12: HR-TEM and SAED images in tibia A) GP-epiphysis interface, from poorly crystalline HAP into plate-like HAp, scale bar = 10 nm; and B) GP-metaphysis interface, from ACP-like structures to poorly crystalline HAp and plate-like HAp, scale bar = 10 nm.

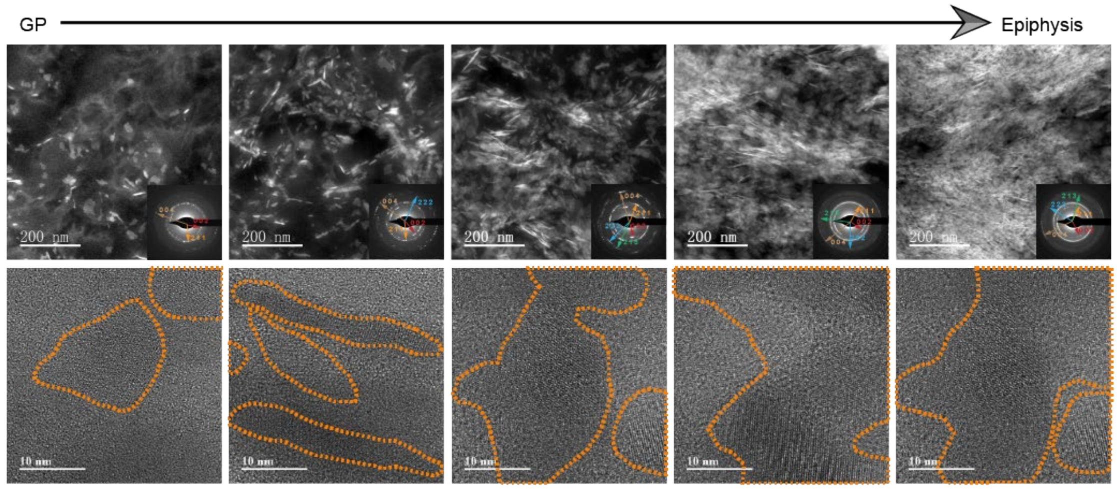

Figure S13: STEM, SAED and HR-TEM images in GP-epiphysis interface region of human phalange tissue, starting from nanocrystals to lacy-like HAp, scale bar = 200 nm in STEM images and 10 nm in HR-TEM images.

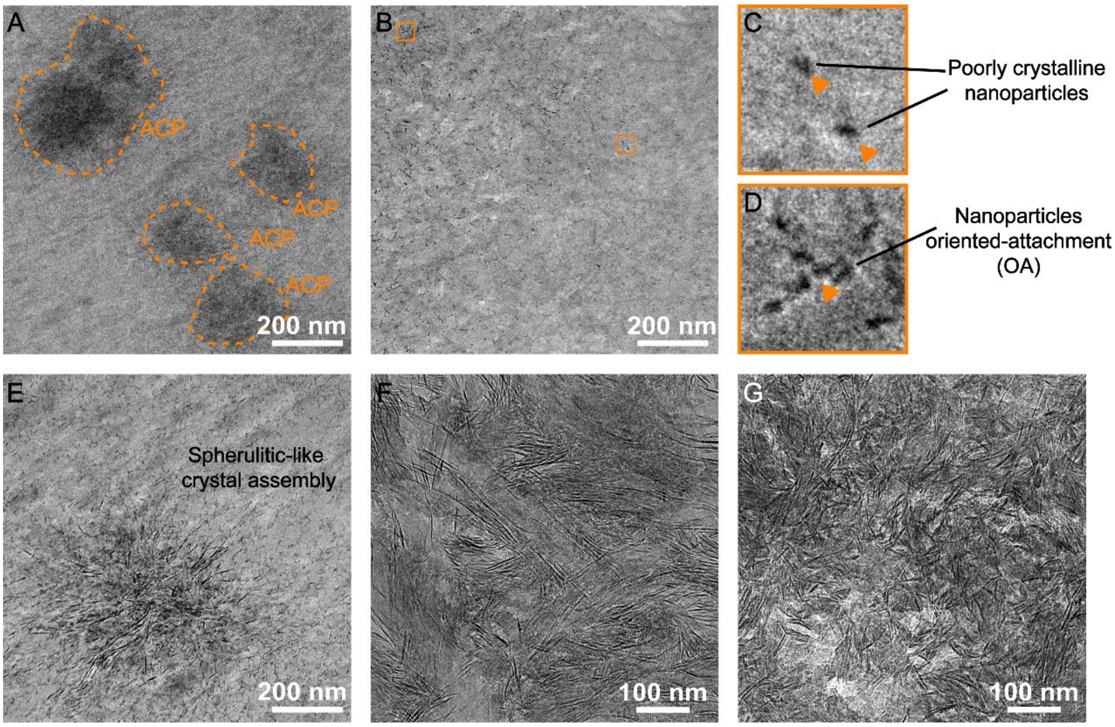


Figure S14: Representative TEM images of tibia GP sample showing the ACP-like structures A), poorly crystalline nanoparticles B, C), nanoparticle oriented-attachment (OA) D), spherulitic-like crystal assembly E) and mineralized region F, G) during GP-metaphysis mineralization process, scale bar = 200 nm in A-E) and 100 nm in F, G).

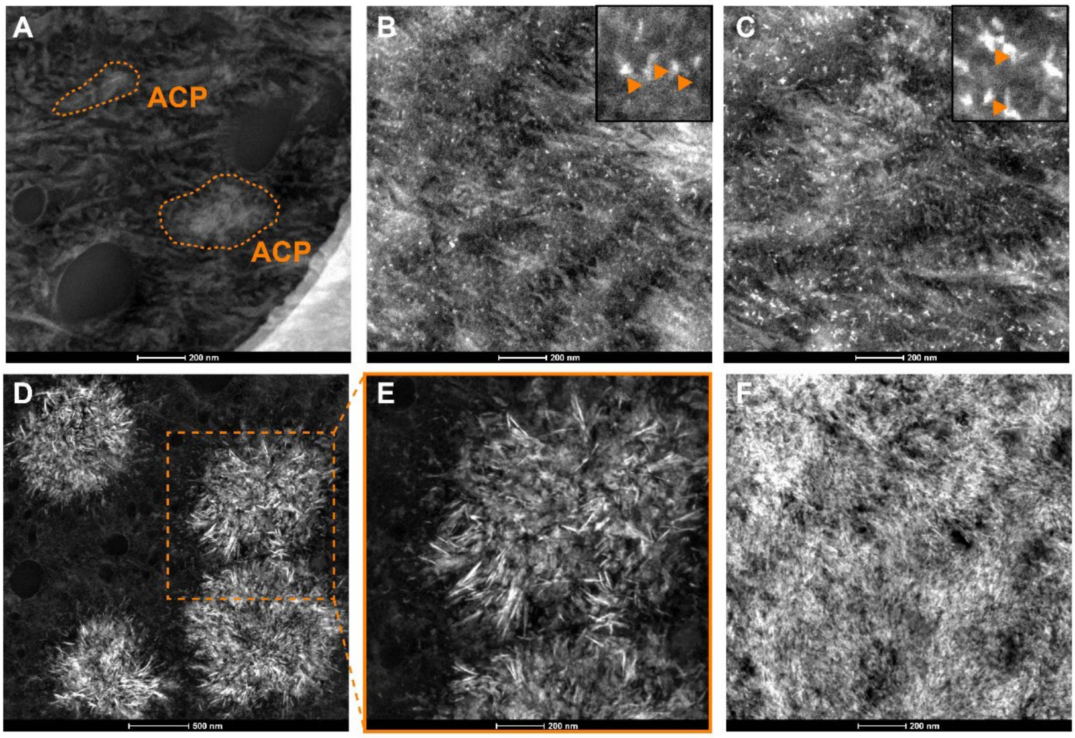

Figure S15: Representative STEM images of tibia GP sample showing the mineralization process of GP-metaphysis interface from A) ACP-like structures to B-C) poorly crystalline nanoparticles, and forming D-E) spherulites and transforming into F) fully mineralized metaphysis tissue, scale bar = 500 nm in D) and 200 nm in A-C, E-F).

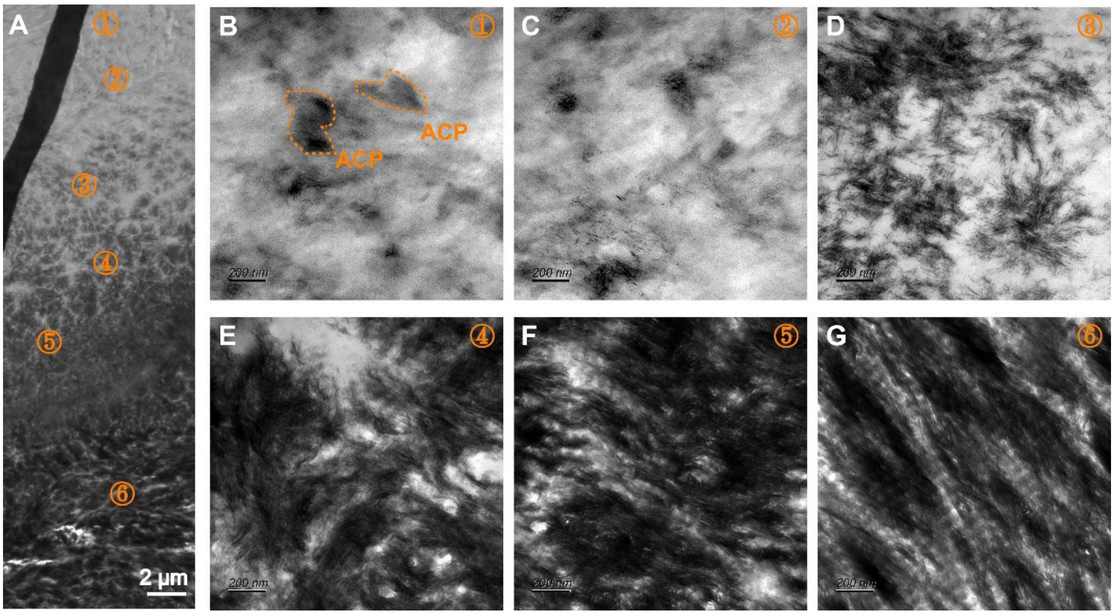


Figure S16: Representative TEM images of GP-metaphysis interface in phalange GP sample showing the gradual mineralization of GP and ACP-like structures B) and spherulitic-like structures D) during biomineralization, scale bar = 2 μm in A) and 200 nm in B-G.

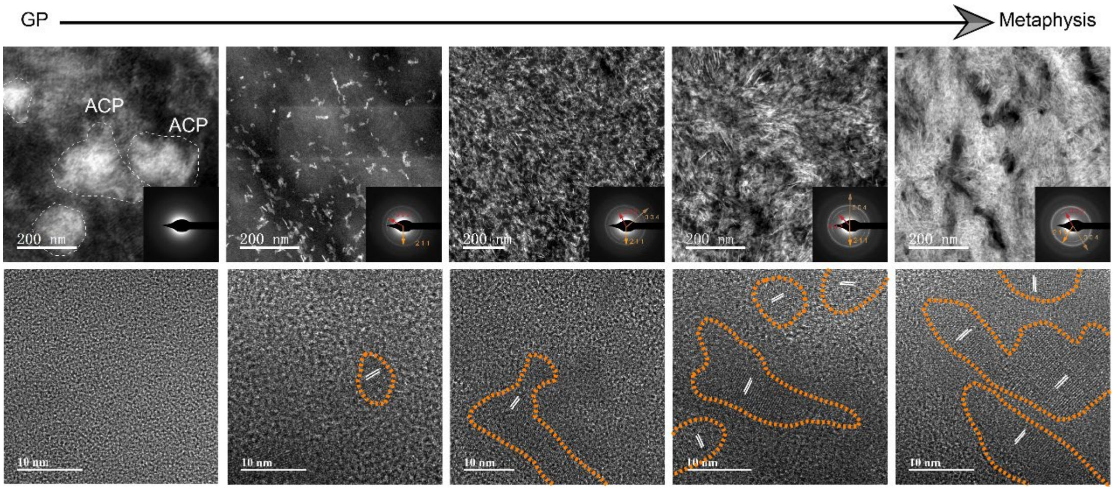


Figure S17: STEM, SAED and HR-TEM images in GP-metaphysis interface region of human phalange tissue, showing the transformation from ACP-like structures to lacy-like HAp, scale bar = 200 nm in STEM images and 10 nm in HR-TEM images.

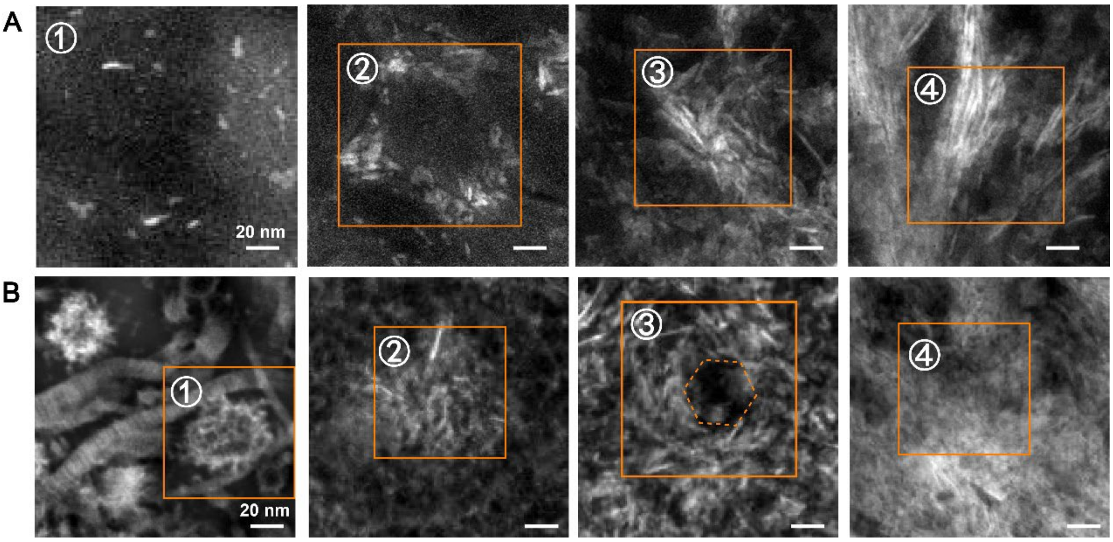


Figure S18: A-B) Representative HHADF-STEM images of minerals from GP-epiphysis A) and metaphysis B), scale bar = 20 nm.

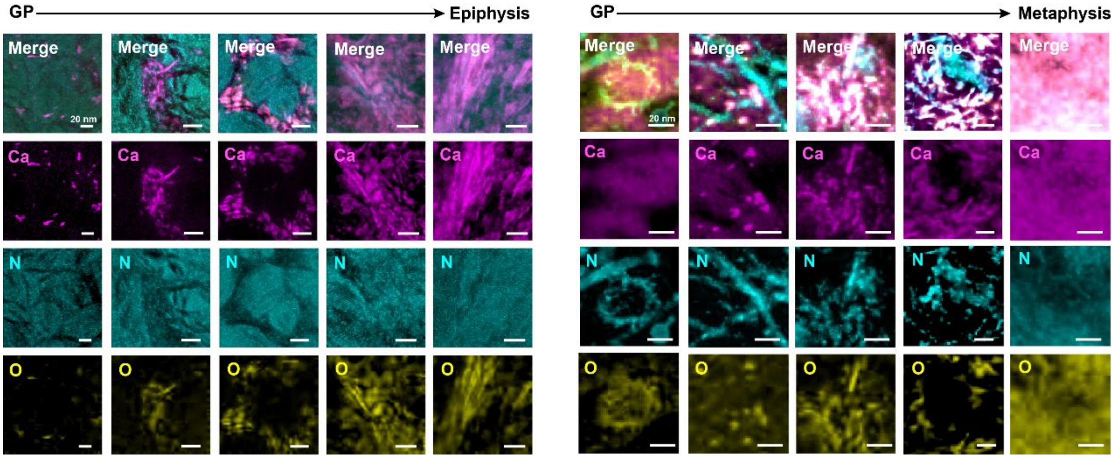


Figure S19: EELS maps corresponding to the spatial distribution of Ca in magenta (collected at Ca L23-edge), O in yellow (collected at O K-edge) and N in cyan (collected at N K-edge) in GP-epiphysis interface and GP-metaphysis interface region, scale bar = 20 nm.

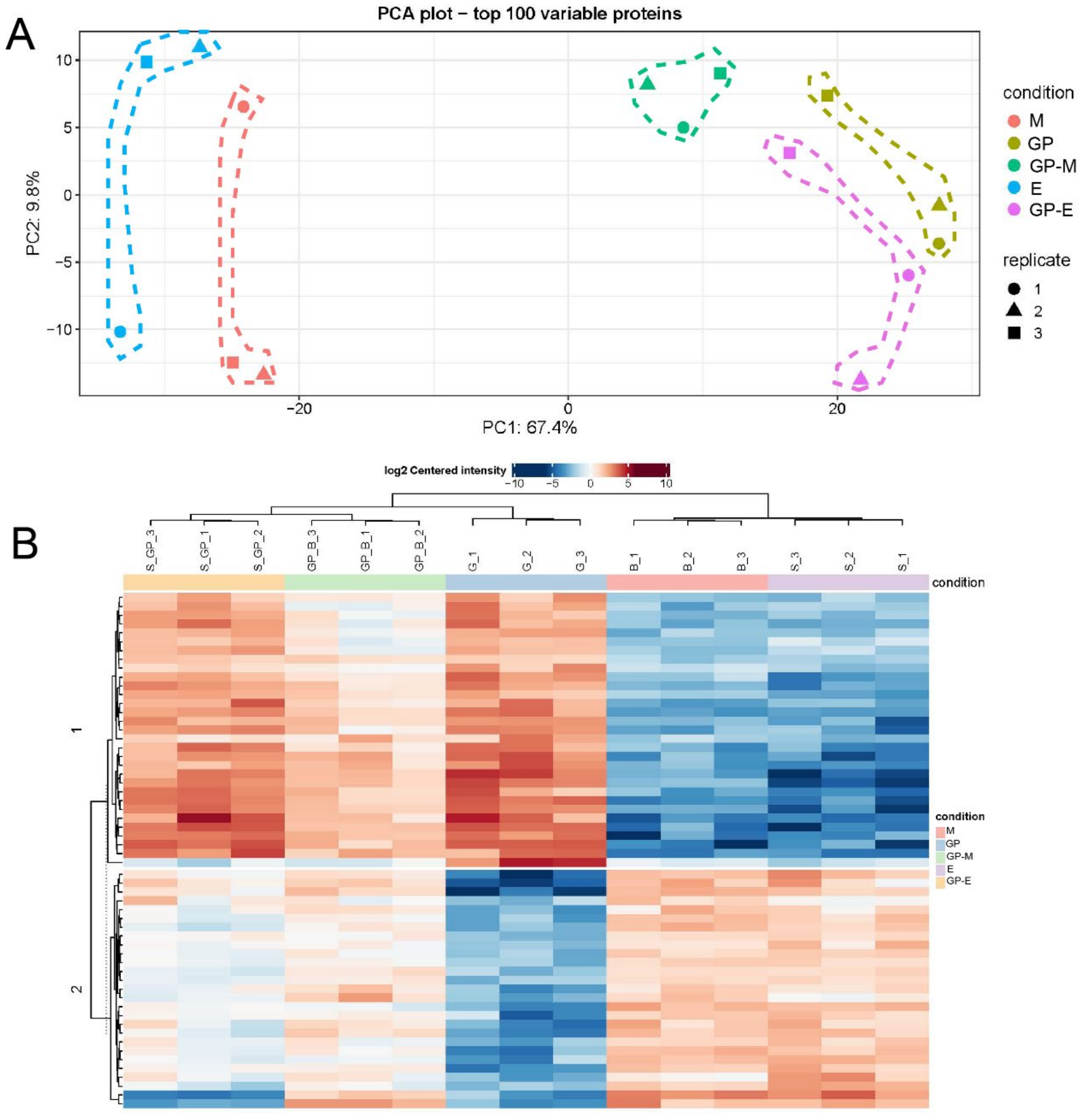


Figure S20: A) The principal component analysis (PCA) of proteomics data and B) heatmap view of proteins identified by proteomics in five groups: epiphysis (E), GP-epiphysis (GP-E), GP, GP-metaphysis (GP-M) and metaphysis (M) tissue.

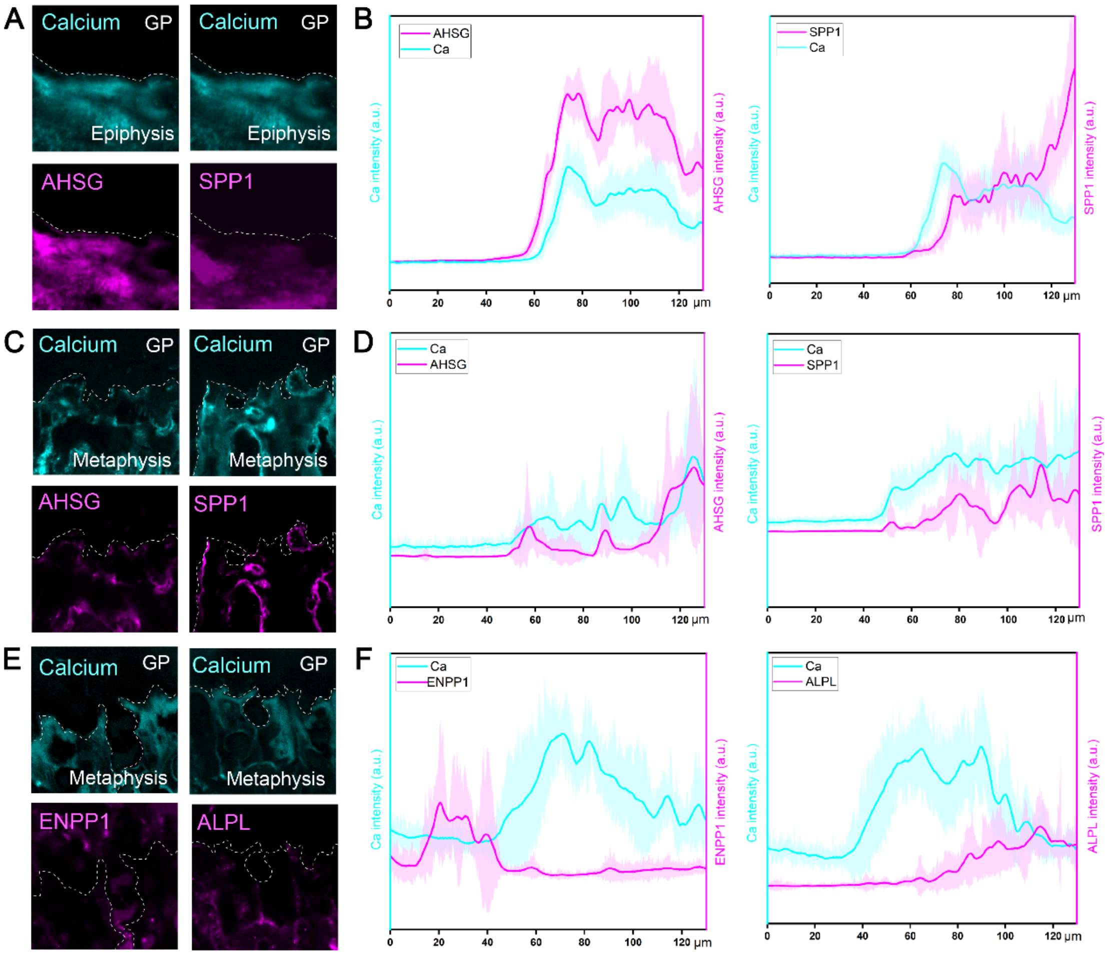


Figure S21: A-B) Representative images of GP-epiphysis interface samples immunostained for AHSG/calcium and SPP1/calcium and corresponding fluorescence intensities plotted over distance across the interface from GP to epiphysis; C-F) Representative images of GP-metaphysis interface samples immunostained for AHSG/calcium, SPP1/calcium, ENPP1/calcium and ALPL/calcium and corresponding fluorescence intensities plotted over distance across the interface from GP to metaphysis, scale bar = 20 µm.

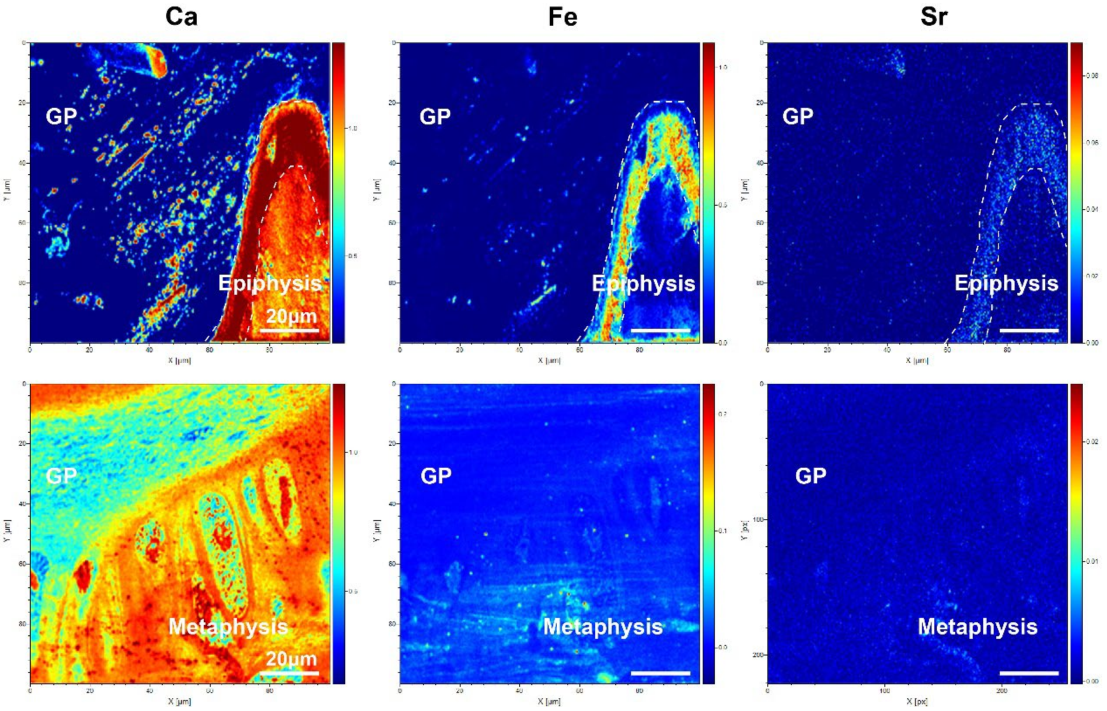


Figure S22: TOF-SIMS images of Ca, Fe and Sr in the region of GP-epiphysis and GP-metaphysis interface, scale bar = 20 μm.

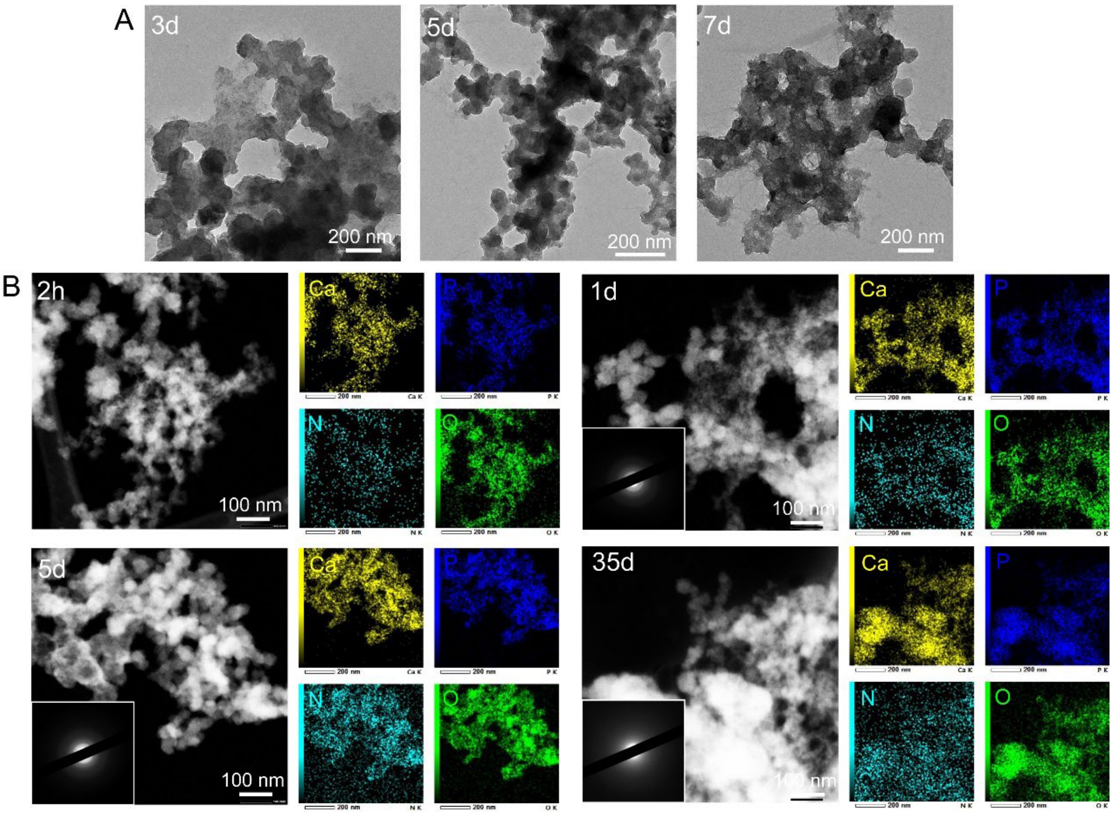


Figure S23: A) Representative cryo-TEM images of prepared ACP after 3d, 5d and 7d at 37 ºC, scale bar = 200 nm; B-D) Representative images of STEM images and EDS mapping images of Ca, P, N, O of prepared ACP particles at 37 ºC after 2h, 1d, 5d and 28d of reaction, scale bar = 100 nm.

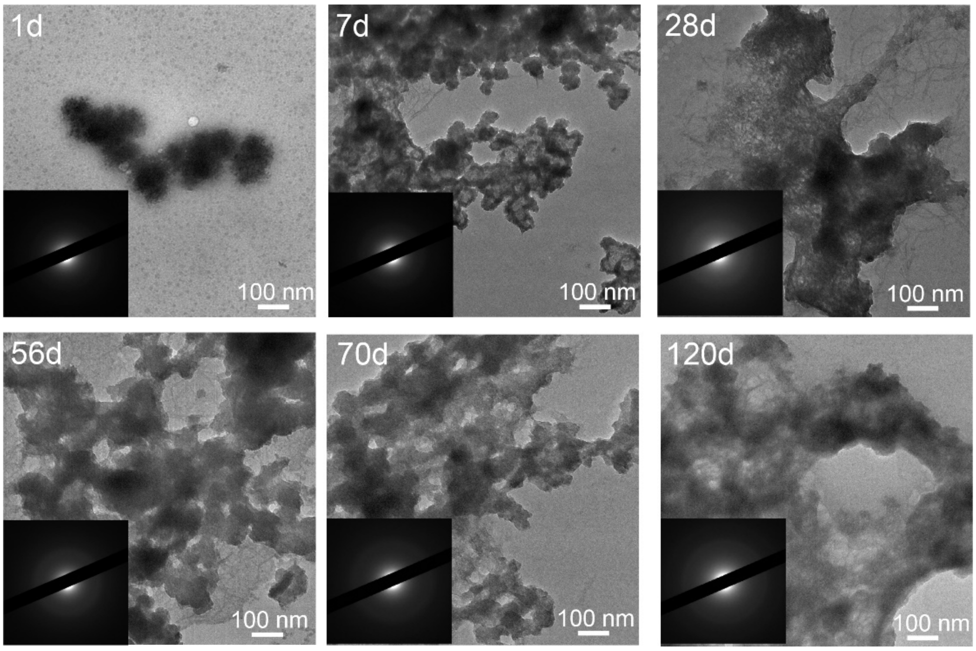


Figure S24: Representative TEM and SAED images of prepared ACP after 1d, 7d, 28d, 56d, 70d and 120d at 4 ºC, scale bar = 100 nm.

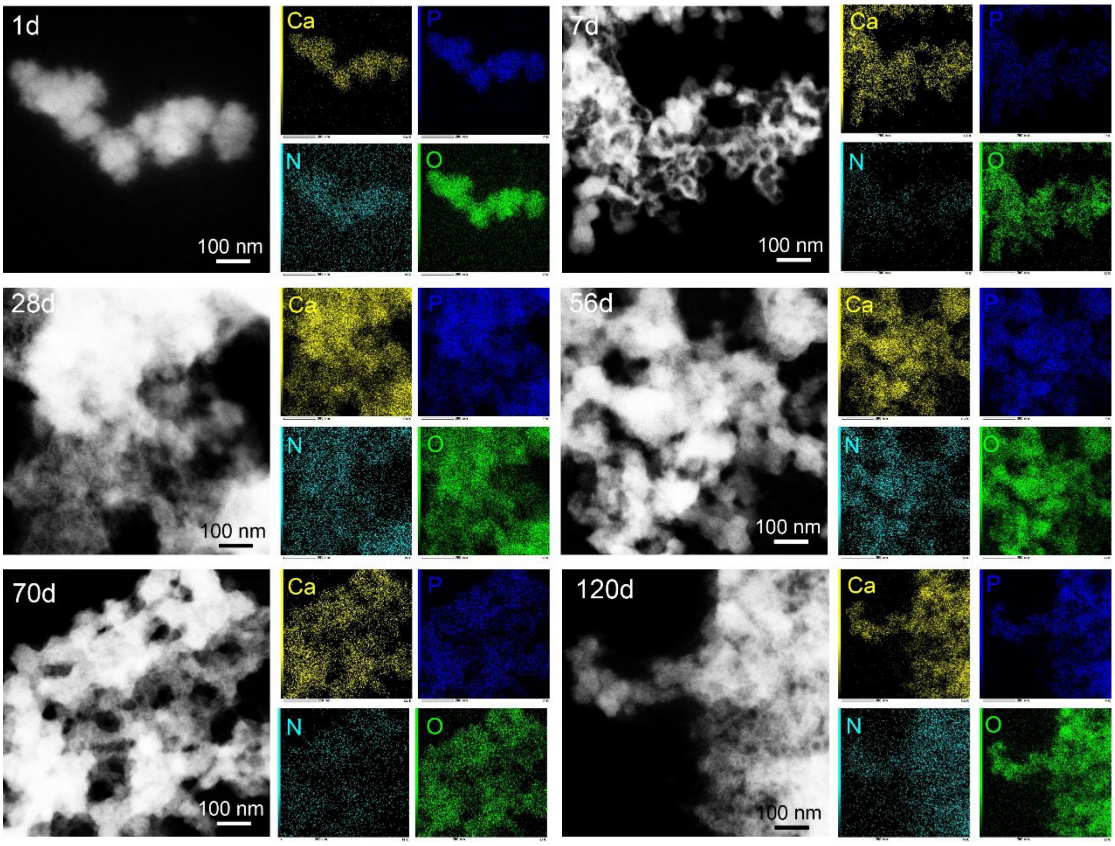


Figure S25: Representative images of STEM images and EDS mapping images of Ca, P, N, O of prepared ACP particles after 1d, 7d, 28d, 56d, 70d and 120d of reaction at 4 ºC, scale bar = 100 nm.

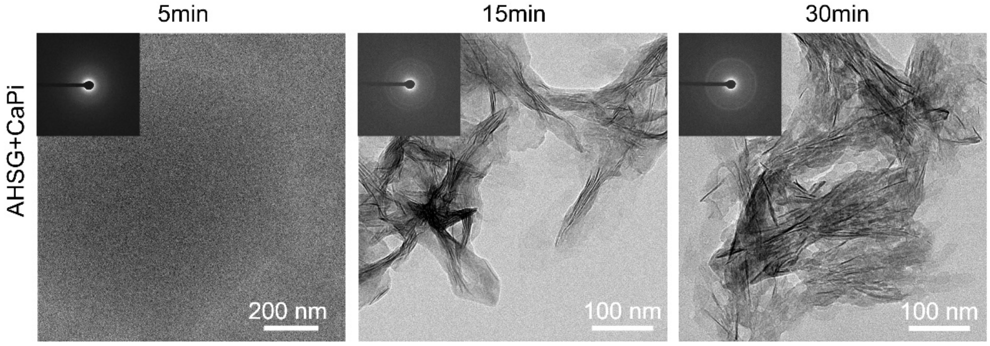


Figure S26: Representative cryo-TEM images of the evolution of structures of AHSG-stabilized calciprotein particles after 5min, 15min and 30min of reaction at 37 ºC, scale bar = 200nm/100nm.

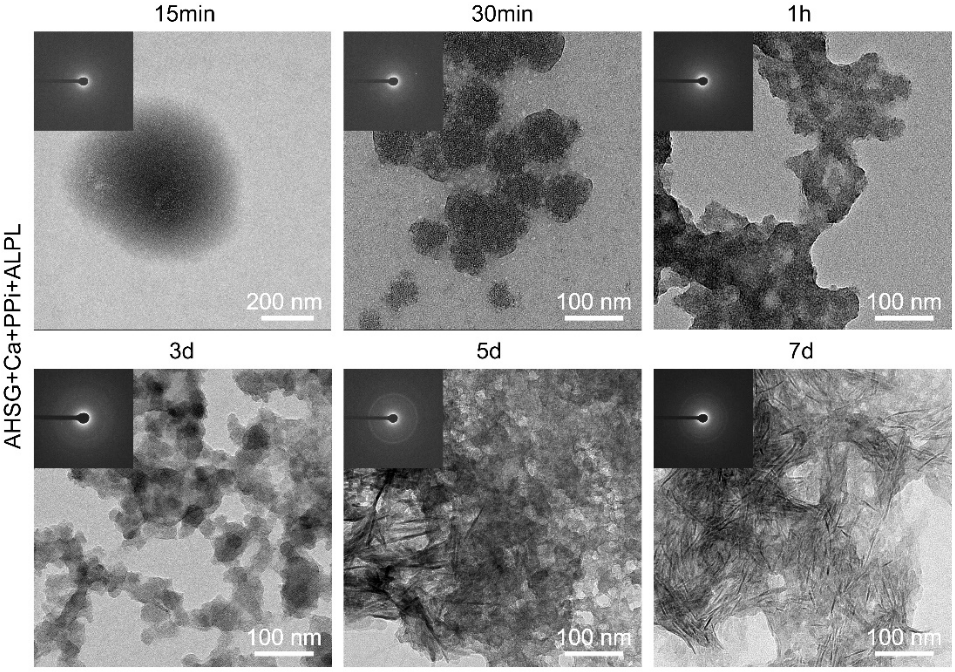


Figure S27: Representative cryo-TEM images of the evolution of structures of AHGS/ALPL-stabilized ACP particles after 15min, 30min, 1h, 3d, 5d and 7d of reaction at 37 ºC, scale bar = 200nm/100nm.

Table S1: Sample information

| Sample source | Numbering scheme | Age | Gender |
|---|---|---|---|
| Phalange samples | Sample 1 | 8m | Male |
| | Sample 2 | 1y7m | Male |
| | Sample 3 | 2y3m | Female |
| | Sample 4 | 5y4m | Female |
| Tibia samples | Sample 1 | 6y | Female |
| | Sample 2 | 6y | Male |
| | Sample 3 | 10y | Male |

| | | | |
|---|---|---|---|
| | Sample 4 | 10y | Female |
| | Sample 5 | 13y | Male |
| | Sample 6 | 14y | Male |

Table S2: EELS characteristic features for minerals, organic compounds and resin

| Phosphorus | Edge | Approximate energy loss | Description |
|---|---|---|---|
| A | P $L_{23}$-edge | 138.0 eV | Transitions to p-like states |
| B | | 141.0 eV | Transitions from 2p state to new state created from due to interactions with calcium 3d orbital function (Characteristic for calcium-containing minerals) |
| C | | 146.0 eV | Transitions to d-like states |
| D | | 160.0 eV | Multiple scattering and the maximum 2p state cross section |
| Carbon | | | |
| A | C K edge | 284.5 eV | Transitions to vacant π*states in in the carbon-carbon bonding environment (Amorphous Carbon) possibly from the embedding resin |
| B | | 287.0 eV | Transitions to the vacant π*state of carbonyl groups possibly from amino acids. |
| C | | 290.2 eV | Transitions to the vacant π*-A states of $CO_3$ groups |
| D | | 297.0 eV | Transitions to vacant σ* states in the carbon-carbon bonding environment |
| Calcium | | | |
| A | Ca $L_{23}$-edge | 348.0 eV | Transitions from $L_3$(2p3/2) to d-like states |
| B | | 351.0 eV | Transitions from $L_2$(2p1/2) to d-like states |
| Nitrogen | | | |
| A | N K edge | 400.0-401.0 eV | 1s-π* transitions |
| B | | 408.0 eV | 1s-σ* transitions in amino compounds |
| Oxygen | | | |
| A | O K edge | 537.0 eV | Transitions to the vacant π* states of the Ca-O bonding environment |
| B | | 539.0 eV | Transitions to the vacant σ* states of the Ca-O bonding environment |

| | | | |
|---|---|---|---|
| C | | 545.0 eV | Transitions to 4s- and 4p-like states in Ca-O bonds |

Movie S1: The displacement magnitude images via DVC processing of GP tissue under compression in XZ plane.
Movie S2: The displacement magnitude images via DVC processing of GP tissue under compression in YZ plane.
Movie S3: The displacement magnitude images via DVC processing of GP tissue under compression in XY plane.
